# Aging represses lung tumorigenesis and alters tumor suppression

**DOI:** 10.1101/2024.05.28.596319

**Authors:** Emily G. Shuldiner, Saswati Karmakar, Min K. Tsai, Jess D. Hebert, Yuning J. Tang, Laura Andrejka, Minwei Wang, Colin R. Detrick, Hongchen Cai, Rui Tang, Dmitri A. Petrov, Monte M. Winslow

**Affiliations:** Department of Biology, Stanford University, Stanford, CA, USA; Department of Genetics, Stanford University School of Medicine, Stanford, CA, USA; Cancer Biology Program, Stanford University School of Medicine, Stanford, CA, USA; The Chan Zuckerberg BioHub, San Francisco, CA USA; Department of Pathology, Stanford University School of Medicine, Stanford, CA, USA

**Author notes:** Corresponding authors: Dmitri A. Petrov Monte M. Winslow.

## Abstract

Most cancers are diagnosed in persons over the age of sixty, but little is known about how age impacts tumorigenesis. While aging is accompanied by mutation accumulation — widely understood to contribute to cancer risk — it is also associated with numerous other cellular and molecular changes likely to impact tumorigenesis. Moreover, cancer incidence decreases in the oldest part of the population, suggesting that very old age may reduce carcinogenesis. Here we show that aging represses tumor initiation and growth in genetically engineered mouse models of human lung cancer. Moreover, aging dampens the impact of inactivating many, but not all, tumor suppressor genes with the impact of inactivating PTEN, a negative regulator of the PI3K/AKT pathway, weakened to a disproportionate extent. Single-cell transcriptomic analysis revealed that neoplastic cells from tumors in old mice retain many age-related transcriptomic changes, showing that age has an enduring impact that persists through oncogenic transformation. Furthermore, the consequences of PTEN inactivation were strikingly age-dependent, with PTEN deficiency reducing signatures of aging in cancer cells and the tumor microenvironment. Our findings suggest that the relationship between age and lung cancer incidence may reflect an integration of the competing effects of driver mutation accumulation and tumor suppressive effects of aging.

## MAIN

Aging and cancer are complex, interrelated processes^1–3^. Epidemiologically, cancer incidence increases with age but does so non-monotonically, with incidence rates rising exponentially before leveling off and ultimately declining in the oldest part of the population ^4–8^. While the age-related increase has long been attributed to the progressive accumulation of driver mutations^9,10^, the deceleration in cancer incidence and mortality among the oldest remains a puzzle and subject of debate^11–13^. Moreover, cancer and aging share many molecular hallmarks, suggesting that there may be additional mechanistic links between these processes^14,15^. Disentangling the complex interplay between aging and tumorigenesis remains a key conceptual and practical challenge for the field.

Lung cancer is the leading cause of cancer death and is highly age-dependent^7^. Autochthonous tumors in genetically engineered mouse models of lung cancer grow within their natural setting and recapitulate critical features of human tumors^16,17^. These tractable systems are uniquely suited to investigate the effects of aging on tumorigenesis because they allow for the controlled induction of cancer-driver mutations. These models can thus separate the age-related increase in the probability of driver mutations from other potential age-related effects on tumorigenesis and eliminate many of the confounding factors that complicate inference from epidemiological and clinical data.

Here we integrate autochthonous models of human lung cancer with tumor barcoding, somatic CRISPR/Cas9 genome editing^18,19^, and single cell transcriptomic analyses to precisely quantify the effects of aging on tumor initiation, subsequent tumor growth, and the functional landscape of tumor suppression. We find that aging represses lung tumorigenesis and lessens the impact of altering several tumor suppressive pathways. Molecular analyses show that the effects of inactivation of PTEN on both cancer cells and the tumor microenvironment are strongly impacted by age. Our work suggests that aging profoundly shapes tumor biology and supports a model in which cancer incidence patterns reflect the competing effects of pro-tumor driver mutation accumulation and anti-tumor effects of tissue aging.

## RESULTS

### Aging represses oncogenic KRAS-driven lung tumor initiation and growth

To assess the impact of aging on lung tumorigenesis, we initiated tumors by intratracheal delivery of a lentiviral vector encoding Cre-recombinase in young (4-6 month old) and old (20-21 month old) mice with *Kras^LSL-G12D/+^* and *Rosa26^LSL-Tomato^* alleles (**Fig. 1a**, **Extended Data Fig. 1a**)^20,21^. The ages of the young and old mice were chosen to correspond to early adulthood and the age at which most molecular phenotypes of aging emerge, respectively^22,23^. Fifteen weeks after tumor initiation, old mice had roughly three-fold lower tumor burden as measured by lung weight and fluorescence imaging (two-sided Wilcoxon rank-sum tests, p=0.028 and 0.0040; **Fig. 1b-d**). There was no difference in lung weight between age-matched non-transduced young and old controls, confirming that weight-based metrics of tumor burden are not confounded by changes with age unrelated to tumor growth (two-sided Wilcoxon rank-sum test, p=0.49; **Extended Data Fig. 1b**) To determine whether reduced efficiency of lentiviral transduction of old lung epithelial cells could explain these results, we transduced young and old wild-type mice with a Lenti-*GFP* vector followed by flow cytometry-based quantification of the percentage of transduced epithelial cells (**Extended Data Fig. 2a**). Transduction efficiency was similar in young and old mice across three replicate experiments, demonstrating that the reduction in tumorigenesis with age is not driven by reduced lung epithelial cell transduction (**Fig. 1e** and **Extended Data Fig. 2b**).

**Fig. 1.**
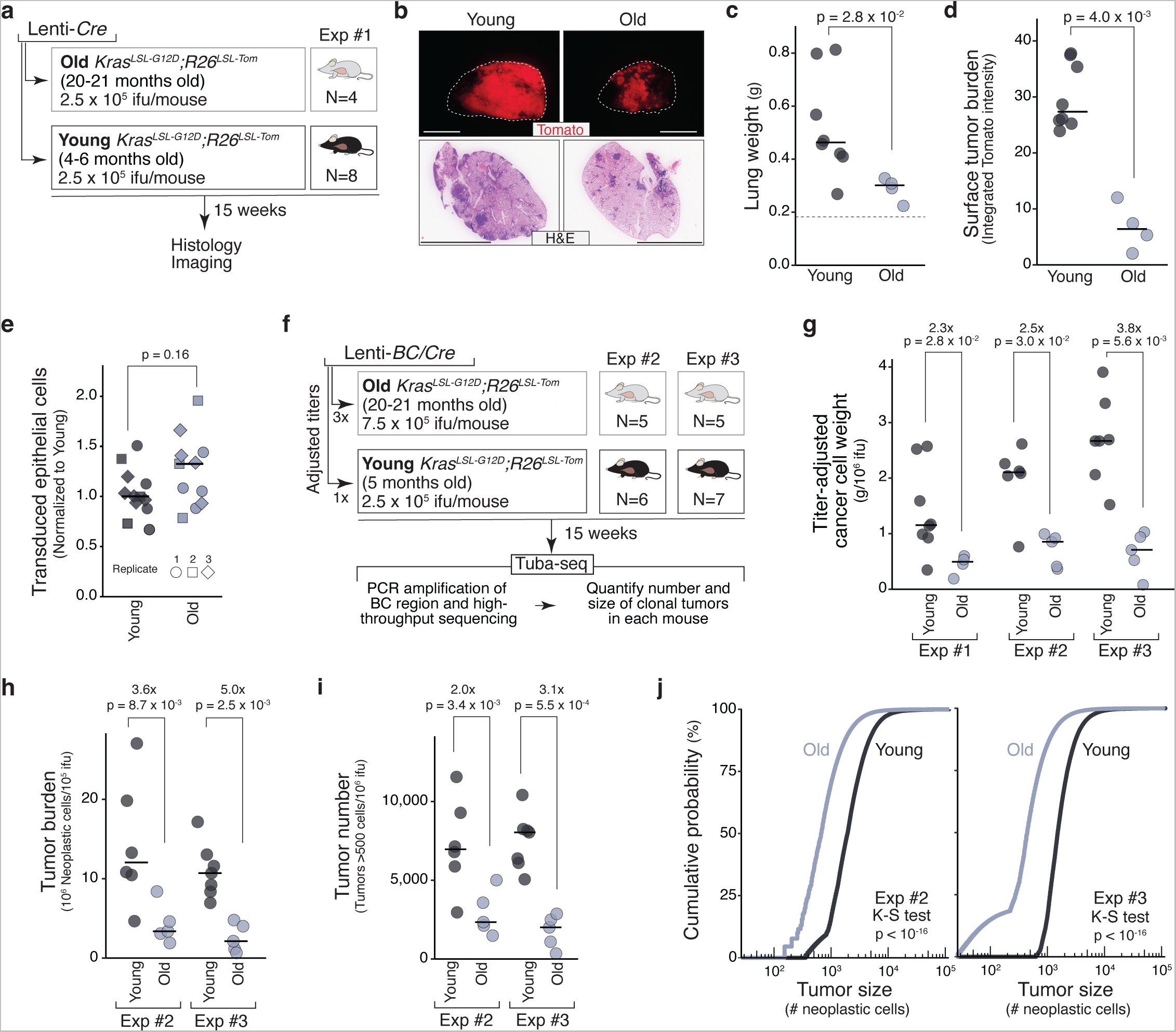
Age reduces oncogenic KRAS-driven lung tumor initiation and growth. **a.** Initiation of KRAS-driven lung tumors in young and old *Kras^LSL-G12D/+^;R26^LSL-Tomato^* mice with Lenti-*Cre* (Experiment #1). Age and number of mice are indicated. See **Extended Data Fig. 1a** for exact ages and sexes. **b.** Representative Tomato fluorescence and H&E images of lungs from young and old mice. Scale bars = 5mm. **c.** Tumor-bearing lung weights of young and old mice. Each dot is a mouse and the bars indicate the median values. Normal lung weight is indicated by the dashed line. **d.** Fluorescence-based quantification of tumor burden in young and old mice. Each dot is a mouse and bars indicate median values. P-value, two-sided Wilcoxon rank sum test. **e.** Quantification of transduction efficiency in young and old mice merged across three replicate experiments. Values are normalized to the median percentage of transduced epithelial cells per experiment. Each point is a mouse; the shapes indicate the replicate experiment. P-value, two-sided Wilcoxon rank sum test. **f.** Initiation of KRAS-driven lung tumors in young and old *Kras^LSL-G12D/+^;R26^LSL-Tomato^*mice using three-fold higher titers of Lenti-*BC/Cre* in the old mice. Two replicate experiments were performed (Experiment #2 and #3). Age and number of mice is indicated. See **Extended Data Fig. 1c** for exact ages and sexes. **g.** Estimated cancer cell weights of young and old mice, normalized to the viral titer delivered. Cancer cell weights are tumor-bearing lung weights minus normal lung weight (median value from **Extended Data Fig. 1b**). Each dot is a mouse and the bars indicate median values. P-values, two-sided Wilcoxon rank sum test. **h,i.** Tumor burden (total number of neoplastic cells in clonal expansions > 500 cells, **h**) and number of tumors (clonal expansions > 500 cells, **i**) in young and old mice in Exp #2 and #3, quantified by Tuba-seq and adjusted for titer. P-values, two-sided Wilcoxon rank sum test. **j.** Empirical cumulative distribution functions of tumor sizes in young and old groups. K-S test: two-sided asymptotic Kolmogorov-Smirnov test comparing the distribution of tumor sizes in young and old mice.

To confirm that age reduces lung tumorigenesis and to generate samples for tumor barcoding coupled with high-throughput barcode sequencing (Tuba-seq)^18,19^, we initiated tumors with a barcoded Lenti-Cre vector (Lenti-*BC*/*Cre*) in two additional groups of young and old mice (**Fig. 1f**, **Extended Data Fig. 1c**). We initiated lung tumors in old mice with a three-fold higher titer of Lenti-*BC*/*Cre* to match the tumor burden in young and old mice and generate material for subsequent molecular analyses (**Fig. 1f**). These old mice had lung weights similar to their young counterparts (**Extended Data Fig. 1d,e**). The magnitude of reduction in tumorigenesis in old mice was consistent across all three experiments, with old mice having a two- to three-fold reduction in tumor burden, when accounting for differences in viral titer (two-sided Wilcoxon rank-sum tests, p = 2.8 × 10^−2^, 3.0 × 10^−2^, 5.6 × 10^−3^; **Fig. 1g**).

To precisely quantify the number of tumors and size of each clonal tumor in each mouse and thereby disentangle the impacts of aging on lung tumor initiation and growth, we extracted DNA from tumor-bearing lungs, amplified the barcode region from the integrated lentiviral vectors, and high-throughput sequenced the amplicon. The number of tumors (number of unique barcoded growths) and the tumor burden (total number of neoplastic cells summed across all tumors) as quantified by Tuba-seq were consistently reduced in old mice. Tumor numbers were reduced two- to three-fold and tumor burdens reduced four- to five-fold. (two-sided Wilcoxon rank-sum test p = 3.4 × 10^−3^, 5.5 × 10^−4^ for tumor number and p = 8.7 × 10^−3^, 2.5 × 10^−3^ for tumor burden; **Fig. 1h,i**). Importantly, tumors were also significantly smaller in old mice, suggesting that aging represses both lung tumor initiation and subsequent tumor growth (K-S test p < 10^−16^; **Fig. 1j**). Tumor burden, number and size were consistently reduced with age in both males and females (**Extended Data Fig. 3a-f**).

### Impacts of tumor suppressor gene inactivation change with age

Alterations in tumor suppressor genes are ubiquitous in cancer and impact cellular processes that are critical to both cancer development and aging^24–27^. Integration of multiplexed CRISPR-based somatic genome engineering and tumor barcoding enables precise quantification of the growth of many genotypes of tumors in parallel^18,19,28^. To quantify the impact of aging on tumor suppressor gene function, we generated a pool of barcoded lentiviral vectors encoding *Cre* and targeting 25 known or putative tumor suppressor genes as well as “sg*Inert*” non-targeting control vectors and a sgRNA targeting an essential gene (*Pcna*) (Lenti-sg*RNA^Aging^/Cre*, **Fig. 2a**)^19^. The tumor suppressor genes included those with strong functional effects on initiation and/or growth in young mice^18,19^ and genes that are frequently mutated in human lung tumors^24,27,29^. We initiated lung tumors with Lenti-sg*RNA^Aging^/Cre* in young and old *Kras^LSL-G12D^*(*K*)*;H11^LSL-Cas9^* and Cas9-negative *K* mice and performed Tuba-seq on tumor-bearing lungs to quantify the size of each tumor of each genotype. (**Fig. 2b**, **Extended Data Fig. 4a,b**). Analysis of all tumors as well as just those initiated with sg*Inert* vectors (which are driven solely by oncogenic KRAS) confirmed that tumor burden, tumor number, and size were all reduced in the old mice (**Extended Data Fig. 5a-e**). Principal component analysis of the number and size of tumors with each sgRNA relative to those with inert sgRNAs distinguished young and old mice, suggesting that aging alters the spectrum of tumor-suppressive effects consistently across mice (**Fig. 2c**).

**Fig. 2.**
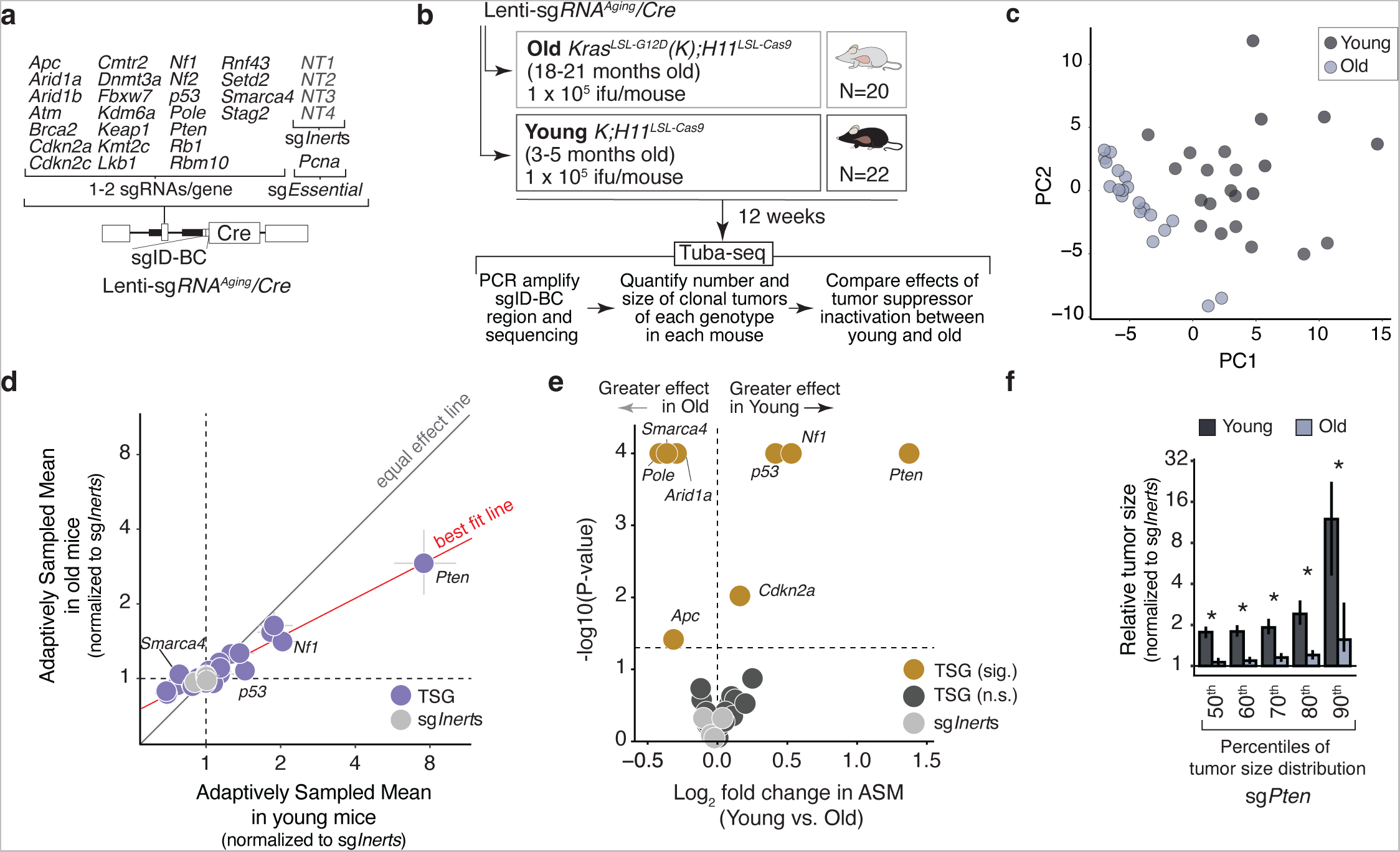
Age alters the landscape of tumor suppression. **a.** Barcoded Lenti-sg*RNA/Cre* vectors in the Lenti-sg*RNA^Aging^/Cre* pool included 2 vectors targeting each putative tumor suppressor, 4 vectors encoding inert (non-targeting) sgRNAs, and a vector with an sgRNA targeting an essential gene. **b.** Initiation of lung tumors in cohorts of young and old *K;H11^LSL-Cas9^*mice with Lenti-sg*RNA^Aging^/Cre*. Ages, genotype, lentiviral titer, and mouse number are indicated. Mice were analyzed after 12 weeks of tumor growth. See **Extended Data Fig. 3a** for exact ages and sexes. **c.** Principal components analysis of tumor suppressive effects calculated within each mouse reveals separation of young and old mice. Each dot is a mouse. **d.** Adaptively sampled mean size (ASM) of each tumor genotype normalized to the ASM of sg*Inert* tumors in the young (x-axis) versus old (y-axis) *K;H11^LSL-Cas9^*mice cohorts transduced with Lenti-sg*RNA^Aging^/Cre*. ASM is a summary metric of tumor fitness that integrates the impact of inactivating each gene on tumor size and number (**Methods**). Confidence intervals were calculated by nested bootstrapped resampling within the young and old cohorts. “Equal effect line”: y=x. “Best fit line”: regression of old ASM onto young ASM for all tumor suppressor genes. TSG: tumor suppressor genes. **e.** Volcano plot showing log2-fold change in the adaptively sampled mean size (ASM) of tumors of each genotype in the young vs old *K;H11^LSL-Cas9^* cohorts. Reported ASMs for each genotype are normalized to the ASM of sg*Inert* tumors. Tumor suppressor genes with a significantly different impact in young and old mice (two-sided FDR-adjusted p-value < 0.05) are highlighted in gold and labelled. sg*Inert* vectors are highlighted in light gray. P-values were calculated using nested bootstrap resampling. **f.** Adaptively sampled sizes of tumors initiated with sg*Pten* vectors in young and old *K;H11^LSL-Cas9^* mice transduced with the Lenti-sg*RNA^Aging^/Cre* pool at indicated percentiles of the tumor size distribution. Each statistic is normalized to the tumor size at the corresponding percentile of the sg*Inert* distribution. Stars denote a statistically significant difference (two-sided FDR-adjusted p-value < 0.05) between young and old. P-values and 95% confidence intervals were calculated using nested bootstrap resampling.

To quantify changes in the effects of inactivating each tumor suppressor gene with age, we sampled the same number of tumors per infectious unit of virus delivered for each sgRNA in each age group and calculated the log-normal mean size of these adaptively sampled tumors relative to sg*Inert* tumors (adaptively sampled mean, ASM; **Methods, Extended Data Fig. 4c**). The effects of tumor suppressor gene inactivation were qualitatively similar in the young and old mice: with the exception of *Smarca4*, which was detrimental to tumor growth only in young mice, the genes that increased tumorigenesis when inactivated in young mice also increased tumorigenesis in old mice (**Fig. 2d**). Interestingly, age reduced the average magnitude of effect of tumor suppressor gene inactivation (p=2.3×10^−14^ for deviation of best fit line from equal effect line), with the impacts of several well-established lung tumor suppressor genes, including *Pten*, *p53*, and *Nf1* significantly reduced in old mice (**Fig. 2d,e**, **Extended Data Fig. 4d**). Conversely, the impacts of inactivating several other canonical tumor suppressor genes, including *Setd2*, *Stag2*, and *Rbm10* were very similar in young and old mice (**Extended Data Fig 4d**). These data suggest that while tumor suppressor function is broadly conserved with age, the molecular changes associated with aging lessen the importance of specific tumor suppressor pathways in constraining tumorigenesis.

Although PTEN suppressed lung tumorigenesis in both young and old mice, *Pten* inactivation increased tumor burden >2.5 times more in young than in old mice (bootstrapped p < 1 × 10^−4^; **Fig. 2e**, **Extended Data Fig. 4d**). This reduction stands out even in the context of the broad dampening of effects with age: in young mice inactivation of *Pten* increased tumorigenesis nearly four times more than inactivation of the next strongest tumor suppressor, while in old mice it was less than twice as strong as the next strongest tumor suppressor (**Extended Data Fig. 4d**). Comparing the size of sg*Pten* and sg*Inert* tumors at various percentiles of the tumor size distribution in young and old mice also consistently showed that age reduces the impact of *Pten* inactivation (**Fig. 2f**). To ensure the robustness of these findings, we repeated our analyses varying the number of tumors sampled per infectious unit of virus, using an alternative statistical method to compare the effects of tumor suppressors between young and old mice (“ScoreRGM”)^30^, and analyzing the impacts of tumor suppressor inactivation separately in males and females. These analyses recapitulated that the effects of most tumor suppressor genes are weakened with age, and consistently found that *Pten* inactivation had the most dramatically age-dependent effect irrespective of sex (**Extended Data Fig. 6**, **Extended Data Fig. 7**).

### Age impacts tumor suppression in P53-deficient lung tumorigenesis

*TP53* is the most frequently mutated tumor suppressor gene in lung adenocarcinoma and is implicated in aging^24,27,29,31,32^. Previous data from young mice suggest that some tumor suppressor genes function differently in the context of p53 deficiency^33^; therefore, we initiated tumors with Lenti-*sgRNA^Aging^/Cre* in young and old *K;p53^flox/flox^;H11^LSL-Cas9^* mice (*KP;H11^LSL-Cas9^*) in which all tumors will be p53-deficient (**Fig. 3a**, **Extended Data Fig. 8a**). After 12 weeks of tumor growth, we performed Tuba-seq on bulk tumor-bearing lungs and compared the adaptively sampled mean tumor sizes for each genotype in young and old mice. This experiment recapitulated several key patterns identified in the *K;H11^LSL-Cas9^* mice, including both the qualitative similarity in tumor-suppressive effects between young and old mice and the broad dampening of the impacts of tumor suppressor gene inactivation in old mice (**Fig. 3b**, **Extended Data Fig. 8b-d**). All tumor suppressors with reduced effects in old *K;H11^LSL-Cas9^*mice also had reduced effects in old *KP;H11^LSL-Cas9^* mice (with the exception of *p53*, which as expected had no impact in either young or old *KP;H11^LSL-Cas9^* mice). *Pten* inactivation again had a strikingly age-dependent effect on tumorigenesis (**Fig 3b,c**, **Extended Data Fig. 8c,d**).

**Fig. 3.**
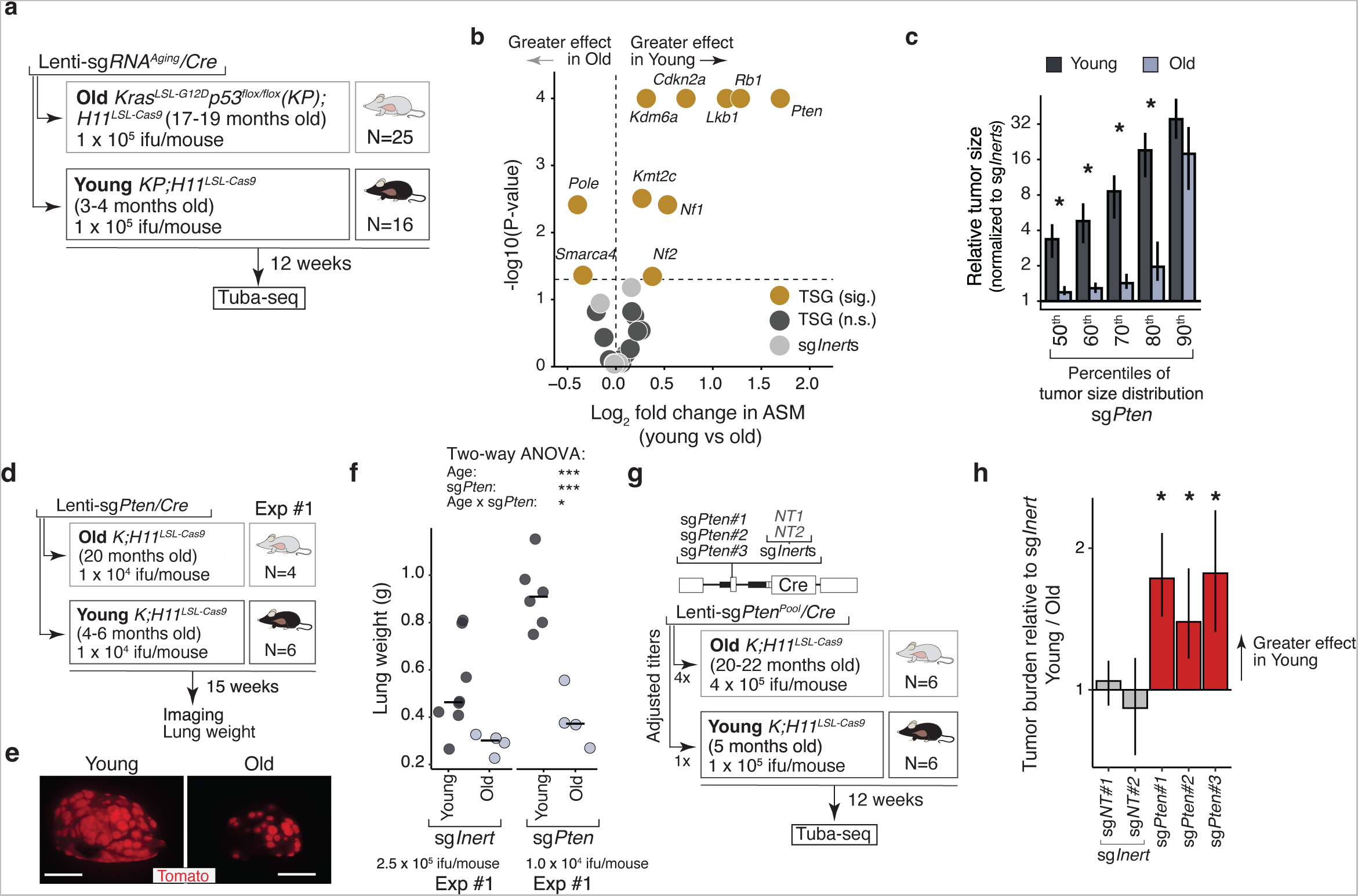
Validation of age-dependent differences in tumor suppressor tumor suppression by *Pten*. **a.** Initiation of lung tumors in young and old *KP;H11^LSL-Cas9^* mice with Lenti-sg*RNA^Aging^/Cre*. Genotype, ages, lentiviral titer, and mouse number are indicated. Mice were analyzed after 12 weeks of tumor growth. See **Extended Data Fig. 8a** for exact ages and sexes. **b.** Volcano plot showing fold change in the adaptively sampled mean size (ASM) of tumors in the young vs old *KP;H11^LSL-Cas9^*cohorts. ASM is a summary metric of tumor fitness that integrates the impact of inactivating each gene on tumor size and number. Reported ASMs for each genotype are normalized to the ASM of sg*Inert* tumors. Tumor suppressor genes with a significantly different impact in young and old mice (two-sided FDR-adjusted p-value<0.05) are highlighted in gold and labelled. sg*Inert* vectors are highlighted in light gray. P-values were calculated using nested bootstrap resampling. **c.** Adaptively sampled percentile sizes of tumors with sg*Pten* vectors in young and old *KP;H11^LSL-Cas9^* mice transduced with Lenti-sg*RNA^Aging^/Cre* pool. Each statistic is normalized to the tumor size at the corresponding percentile of the sg*Inert* distribution. Stars denote a statistically significant difference between young and old (two-sided FDR-corrected p-value<0.05). P-values and 95% confidence intervals were calculated using nested bootstrap resampling. **d.** Schematic of tumor initiation in young and old mice with Lenti-sg*Pten/Cre*. See **Extended Data Fig. 8e** for exact ages and sexes. **e.** Representative Tomato fluorescence images of lungs of young and old mice with tumor initiated with Lenti-sg*Pten/Cre*. Scale bars = 5mm. **f.** Lung weights of young and old mice transduced with Lenti-sg*Pten/Cre*. Two-way ANOVA indicates that there is a statistically significant interaction between the effects of age and *Pten* inactivation on lung weight (F(1,18) =4.66, p=0.45). ***: p < 1 × 10^−3^, **: p < 1 × 10^−2^, *: p < 5 × 10-2. **g.** Schematic of tumor initiation in young and old mice with Lenti-sg*Pten^Pool^/Cre*. See **Extended Data Fig. 8e** for exact ages and sexes. **h.** Tumor burden per sgRNA relative to sg*Inert* tumor burden, in young relative to old mice transduced with Lenti-sg*Pten^Pool^/Cre*. Stars denote a significant differential effect with age (two-sided FDR-corrected p-value<0.05). P-values and 95% confidence intervals were calculated using nested bootstrap resampling.

Notably, there was also consistency in which tumor suppressor genes did not interact with age within p53-proficient and p53-deficient tumors. Age did not impact the effects of inactivation of *Stag2, Setd2, Cdkn2c*, and *Rbm10* in either context, further suggesting that age interacts with specific tumor suppressor pathways as opposed to generically weakening the effects of driver mutations (**Fig. 3b**, **Extended Data Fig. 8c**). Sensitivity analyses again showed that these effects are present in both males and females and are robust to variation in the number of tumors sampled and the use of an alternative statistical method (**Extended Data Fig. 6**, **Extended Data Fig. 7**).

### Age weakens the tumor suppressive effect of PTEN

*Pten* inactivation had the greatest reduction in effect with age among the tumor suppressor genes we assayed in both oncogenic KRAS and KRAS;P53-deficient lung tumors. To validate the impact of aging on the tumor-suppressive function of PTEN outside of a pooled setting, we initiated tumors in young and old *Kras^LSL-G12D^;Rosa26^LSL-tdTomato^;H11^LSL-Cas9^*(*KT;H11^LSL-Cas9^*) mice with a lentiviral vector encoding Cre recombinase and a *Pten*-targeting sgRNA (“*sgPten”* vector, **Fig. 3d**, **Extended Data Fig. 8e**). Old mice had visibly fewer tumors and dramatically lower lung weights (**Fig. 3e,f**). Notably, the reduction in lung weight with age in mice transduced with the sg*Pten* vector exceeded that of mice transduced with the sg*Inert* vectors (two-way ANOVA p=0.045 for sg*Pten* by age interaction term; **Fig. 3f**).

To further validate that age reduces the tumor-suppressive function of PTEN and confirm that this result is driven by on-target effects, we generated a pool of barcoded Lenti-sgRNA/Cre vectors containing three distinct sgRNAs targeting *Pten* as well as two sg*Inert* control vectors (Lenti-sg*Pten^Pool^/Cre*). We initiated tumors with Lenti-sg*Pten^Pool^/Cre* in young and old *KT;H11^LSL-Cas9^* mice and performed Tuba-seq on bulk-tumor bearing lungs (**Fig. 3g**, **Extended Data Fig. 8e**). We calculated the impact of each sgRNA on tumor burden relative to the sg*Inert* vectors and compared this metric between young and old mice. In addition, we calculated scoreRGM^30^ for each sgRNA. All three *Pten*-targeting sgRNAs had a greater impact on tumor burden in young mice than in old mice by both metrics (**Fig. 3h**, **Extended Data Fig. 8f**). Thus, the tumor-suppressive effect of *Pten* is weakened with age across three distinct sgRNAs and across four separate experiments on young and old mice.

### Age has a continued impact on the gene expression state of neoplastic cells

To assess changes in neoplastic cell state and the tumor microenvironment with age and gain insights into the age-dependent effect of PTEN, we performed single-cell RNA-sequencing (scRNA-seq) on cells from the tumor-bearing lungs of young and old *KT;H11^LSL-Cas9^* mice with tumors initiated with Lenti-sg*Inert/Cre* (sg*Inert*) and Lenti-sg*Pten/Cre* (sg*Pten*) (n=4 mice per age-genotype group; **Fig. 4a**)(**Extended Data Fig. 9a,b**). We separately enriched for neoplastic cells and for all other stromal and immune cells by FACS (**Fig. 4a, Methods**). After quality controls including *in silico* removal of transduced non-neoplastic cells, ~180,000 cells (~100,000 from young and ~80,000 from old mice) were assigned to 19 expected cell types in the mouse lung (**Fig. 4b-c**, **Extended Data Fig. 9c,d, Methods**)^34,35^. Each cell type included cells from each age group and genotype, indicating that neither age nor *Pten* inactivation confounded cell type annotation (**Fig. 4c**, **Extended Data Fig. 9e-i**). Across the five major epithelial-like cell subtypes, *Tomato*-expressing cells were greatly enriched in alveolar type 2 (AT2), AT1/2- and AT1-like populations. We therefore focused on these cells as the neoplastic/cancer cell fraction (**Extended Data Fig. 10a-i, Methods)**.

**Fig. 4.**
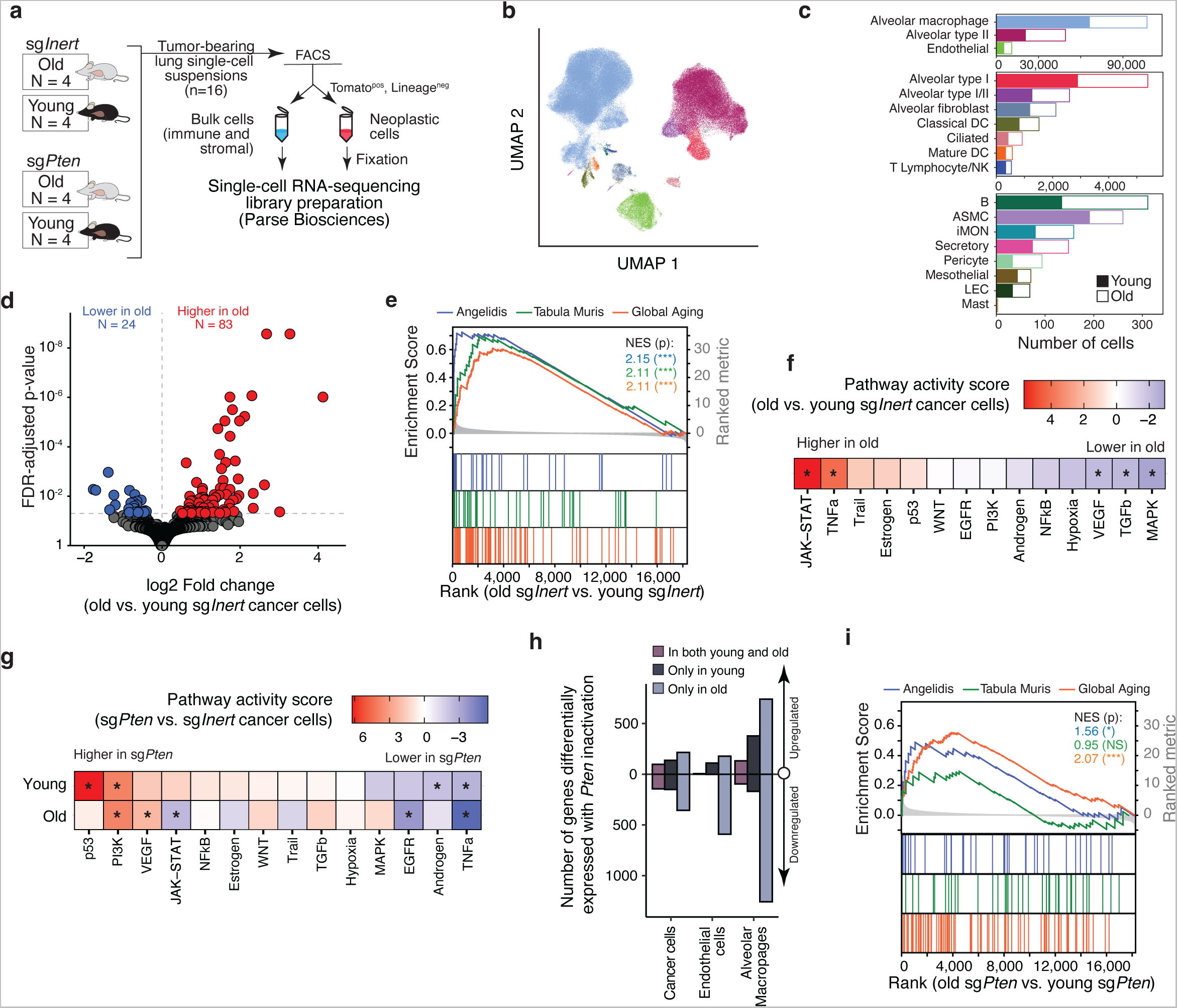
Aging alters cancer cell state and the impacts of *Pten* inactivation. **a.** Single-cell RNA sequencing of cells from tumor-bearing lungs of young and old *KT;H11^LSL-Cas9^* mice transduced with Lenti-sg*Inert/Cre* or Lenti-sg*Pten/Cre* vectors. To enrich for cancer cells, whole-lung single-cell suspensions were divided into a cancer-cell fraction and a bulk fraction containing all other live cells by fluorescence-activated cell sorting (FACS). Lineage: markers of non-epithelial lineages (CD45, CD31, Ter119, F4/80). **b.** Uniform Manifold Approximation and Projection (UMAP) embedding of cells colored by cell type. Mapping of colors to cell types is indicated in (**c**). **c.** Counts per cell type by age. DC: dendritic cells, ASMC: airway smooth muscle cells, iMON: inflammatory monocytes, NK: natural killer cells, LEC: lymphatic endothelial cells. **d.** Volcano plot showing changes in gene expression with age in cancer cells from the sg*Inert* samples. Differentially expressed genes (Benjamini-Hochberg corrected Wald test p<0.05) are highlighted in color. **e.** Gene expression signatures of normal aging are enriched with age in KRAS-driven lung cancer cells. Angelidis: genes with positive log fold change with aging and adj. p value <0.05 in Angelidis et al., Tabula Muris: lung type II pneumocyte aging gene set from Tabula Muris Consortium, Global Aging: global aging genes upregulated with age from Zhang et al. NES: Normalized enrichment statistic. P-values calculated by permutation and FDR-corrected. ***: p < 5 × 10^−3^. **f.** Pathway activity scores comparing old to young cancer cells from mice with sg*Inert* tumors. Positive scores (red) indicate that a pathway is more active with age; negative scores (blue) indicate that a pathway is less active with age. Stars denote that a pathway is differentially activated with age (p<0.05, PROGENy multivariate linear model). **g.** Pathway activity scores comparing *Pten* deficient (sg*Pten*) to wildtype (sg*Inert*) for young and old cancer cells. Positive scores (red) indicate that a pathway is more active with *Pten* inactivation; negative scores (blue) indicate that a pathway is less active with *Pten* inactivation. Stars denote that a pathway is differentially activated with age (p<0.05, PROGENy multivariate linear model). **h.** Numbers of genes differentially expressed with *Pten* inactivation in indicated cell types, stratified by whether genes were up/down regulated, and by whether effect is specific to young or old cells, or is common to both ages. **i.** Enrichment of gene expression signatures of normal aging (as in **e**) are weakened in *Pten*-deficient lung tumors. ***: p < 5 × 10^−3^, *: p < 5 × 10^−2^. NS: not significant.

Single-cell mapping efforts have revealed extensive transcriptomic changes with age^23,36–39^. Oncogenic transformation is a major departure from cellular homeostasis^15^. Thus, it is unclear whether cells in the tumors of old mice will retain these canonical features of aging or will converge to a shared molecular state with young cancer cells. sg*Inert* neoplastic cells from tumors in young and old mice were represented similarly within the major epithelial-like cell clusters, suggesting that age does not lead to dramatic shifts in cell identity (**Extended Data Fig. 10i**). However, pseudo-bulk comparison between young and old neoplastic cells uncovered significantly differently expressed genes (**Fig. 4d**). Neoplastic cells from tumors that developed in old mice (“old cancer cells”) had a striking enrichment for several signatures of lung epithelial^23,36^ and global aging^40^ when compared to neoplastic cells from tumors that developed in young mice (“young cancer cells”), suggesting that oncogenic KRAS-driven lung tumorigenesis does not revert these fundamental changes (**Fig. 4e**).

To further understand the impacts of age on the molecular state of cancer cells, we used PROGENy^41^ to infer changes in the activation of canonical signaling pathways with age. Interestingly, several pathways that could logically impact tumorigenesis were differentially active with age, including reduced MAPK signaling in old cancer cells, which has a critical role in driving proliferation downstream of oncogenic KRAS (**Fig. 4f**). Thus, the impact of age on KRAS-driven lung tumors continues beyond initiation and the earliest stage of tumorigenesis.

### Age changes the transcriptomic effects of *Pten* inactivation on cancer cells and the tumor microenvironment

*Pten* inactivation has a highly age-dependent impact on tumor fitness (**Fig. 2,3**). To gain insight into the molecular basis of this genotype-by-age interaction, we compared the effects of *Pten* inactivation on the transcriptomes of young and old cancer cells. Consistent with the function of PTEN as a canonical negative regulator of PI3K signaling^42^, this pathway was upregulated to a similar extent in both young and old sg*Pten* cancer cells. However, changes in the activation of several other pathways were specific to a single age context (**Fig. 4g**). These results argue against a simple model in which age reduces the fitness benefit associated with *Pten* inactivation through curbing activation of the pathway that it suppresses, but instead suggest that aging alters the molecular consequences further downstream of PI3K activation.

The frequencies of most immune and stromal cell types were unchanged with either age or *Pten* inactivation (**Extended Data Fig. 11a**). However, the growth of PTEN deficient tumors strongly impacted the transcriptional state of stromal and immune cells, with particularly strong effects on alveolar macrophages and endothelial cells (**Extended Data Fig 11b**). Interestingly, these effects were also highly age-dependent: for each cell type, the majority of genes that were up- or down-regulated with *Pten* inactivation did so specifically in young or old cells (**Fig. 4h**, **Extended Data Fig 11c**). Thus, aging not only shapes the consequences of PTEN inactivation on the molecular state of cancer cells, but also alters non-cell autonomous effects of the growth of PTEN-deficient tumors.

### *Pten*-deficiency reduces molecular phenotypes of aging

To better understand the age-specific consequences of PTEN inactivation, we directly compared the transcriptomes of young and old sg*Pten* cancer cells. Strikingly, young and old sg*Pten* cancer cells had fewer differences in gene expression, muted changes in signaling pathway activation, and weaker enrichment of aging signatures relative to young and old sg*Inert* cancer cells (**Fig. 4i**, **Extended Data Fig. 12a,b**). Likewise, per-cell aging scores of both young and old sg*Pten* cancer cells resembled those of young sg*Inert* cancer cells, suggesting that PTEN inactivation in old cancer cells may drive transcriptomic changes that oppose aging phenotypes (**Extended Data Fig. 12c**). Consistent with this, the impacts of *Pten* inactivation in old cancer cells were negatively correlated with the effects of aging at the pathway level (r = −0.65), and roughly a third of the genes differentially expressed with age were regulated in the opposite direction with *Pten* inactivation in old cancer cells (**Extended Data Fig. 12d-e**).

Finally, the effects of *Pten* inactivation extended to cells in the tumor microenvironment. Age-associated transcriptional changes in stromal and immune cells from mice with sg*Inert* tumors were almost entirely absent in mice with sg*Pten* tumors (**Extended Data Fig. 12f-h**), and most cell types in young and old mice with sg*Pten* tumors were transcriptomically as young or younger than the corresponding cells in mice with sg*Inert* tumors (**Extended Data Fig. 12h**). These results suggest that *Pten* inactivation restores a youthful state in old tumors and surrounding tissue, highlighting that the phenotypic consequences of driver mutations can both impact and be impacted by aging.

## DISCUSSION

Understanding how aging impacts tumorigenesis is a key challenge for the field which grows more pressing as the global population ages^43,44^. Epidemiological studies have suggested a protective effect of very old age, but interpretation of these data are complicated by age-associated tumor-extrinsic factors including differences in diagnostic intensity, patterns of environmental exposure, and the prevalence of co-morbidities^11,45^. Here we used genetically engineered mouse models of human lung cancer to characterize the effects of age on tumorigenesis independently of age-associated driver mutation accumulation. We find that aging reduces tumor initiation and growth driven by oncogenic KRAS, one of the most common oncogenic drivers across all human cancers. While reduced tumor initiation may reflect exhaustion of alveolar progenitor cells in the lung with age^46^, slower tumor growth post-initiation suggests that aging also weakens the proliferative ability of transformed cells. Our transcriptomic analysis supports this notion, as old cancer cells fail to converge to a shared phenotypic state with young cancer cells and have reduced MAPK signaling. We also find that KRAS-driven tumors respond differentially to tumor suppressor inactivation with age, with the impacts of several important tumor suppressor genes dampened with age. These findings are consistent with age not only reducing the fitness of KRAS-driven lung tumors, but also constraining their maximum fitness and thereby limiting the beneficial effects of secondary mutations.

This reduction in effect was especially pronounced for PTEN, a canonical negative regulator of the PI3K/AKT pathway, a central oncogenic signaling pathway in many cancer types which is activated through a variety of mechanisms in a large fraction of lung adenocarcinomas^47–49^. Our scRNA-seq analysis showed that while PTEN inactivation in cancer cells strongly upregulated PI3K signaling irrespective of age, the downstream phenotypic effects of this activation shifted dramatically with age in both cancer and stromal cells. These differing effects may indicate a rewiring of the signaling landscape in old cancer cells that ultimately constrains the oncogenic potential of this important signaling axis. Surprisingly, PTEN-deficient tumors as well as co-existing stromal cells from old mice had reduced transcriptomic signatures of aging relative to mice with PTEN-proficient tumors. This finding is unexpected given the lifespan-extending effects of reduced PI3K signaling^50–52^. Collectively, these findings underscore the interconnectedness of pathways involved in aging and tumorigenesis.

Although we focus on KRAS-driven lung adenocarcinoma in this study, aging may be a generic tumor-suppressive mechanism. If so, the non-monotonic pattern of cancer incidence with age may reflect an integration of the competing effects of pro-tumor driver mutation accumulation and general anti-tumor tissue aging, where aging-induced tumor-suppressive changes begin to dominate in the very elderly even as somatic cells continue to accrue genomic alterations. Importantly, both the dynamics of mutation accumulation and the tumor-repressive effects of aging are likely tissue-dependent, necessitating further investigation on the impacts of aging in different cancer types and tissue contexts. A key future challenge will be to identify the specific aspects of aging that suppress tumor growth so that they can be harnessed separately from the pathological effects of aging.

## METHODS

### Design and generation of Lenti-sgRNA/Cre vectors

Most barcoded Lenti-sgRNA/Cre vectors have been described (Cai et al). New sgRNA sequences targeting *Pten* were designed using CRISPick (https://portals.broadinstitute.org/gppx/crispick/public). sgRNA sequences are presented in **Supplementary Table 1**. Each desired sgRNA vector was modified from pll3-U6-sgRNA-Pgk-Cre vector via site-directed mutagenesis (New England Biolabs, E0554S). The generation of the barcode (BC) fragment containing the 8-nucleotide sgID sequence and 20-nucleotide degenerate BC, and subsequent ligation into the vectors, were performed as previously described^1,2^.

Lenti-sgRNA/Cre vectors were individually co-transfected into 293T cells with pCMV-VSV-G (Addgene #8454) envelope plasmid and pCMV-dR8.2 dvpr (Addgene #8455) packaging plasmid using polyethylenimine in 150 mm cell culture plates. Sodium butyrate (Sigma-Aldrich, B5887) was added 8 h after transfection to achieve a final concentration of 20 mM. Medium was refreshed 24 h after transfection. Supernatants were collected at 36 and 48 hours after transfection, filtered through a 0.45 μm syringe filter (Millipore SLHP033RB) to remove cells and debris, concentrated by ultracentrifugation (25,000 g for 1.5 hours at 4°C), and resuspended overnight in PBS then frozen at −80 °C.

### Lentiviral titering and pooling

Concentrated lentiviral particles were titered by transducing LSL-YFP cells (a gift from Dr. Alejandro Sweet-Cordero/UCSF), determining the percent YFP-positive cells by flow cytometry, and comparing the titer to a lentiviral preparation of known titer. Vectors for the Tuba-seq studies in *K;H11^LSL-Cas9^* and *KP;H11^LSL-Cas9^* mice were pooled with the goal of producing roughly equal tumor burden per vector in the young mice based on prior Tuba-seq data. Specifically, all vectors were pooled at equal titers with the exception of vectors targeting genes known to be strong tumor suppressors (*Pten*, *Lkb1*, *Nf1*, *Stag2*, and *Setd2*), which were pooled at half titers. For calculations of tumor suppressive effect, the exact representation of each vector in the viral pool was determined through analysis of tumor-bearing lungs from Cas9-negative *K* control mice (see **Adaptive sampling of tumors for statistical comparison of tumor genotypes**).

### Mice, tumor initiation, and tissue collection

The use of mice for this study was approved by the Institutional Animal Care and Use Committee at Stanford University, protocol number 26696. *Kras^LSL-G12D/+^*(RRID:IMSR_JAX:008179), *p53^flox/flox^* (RRID:IMSR_JAX008462), *R26^LSL-tdTomato^* (RRID:IMSR_JAX:007914), and *H11^LSL-Cas9^*(RRID:IMSR_JAX:027632) mice have been previously described^3–6^. All mice were on a *C57BL/6* background. The sex of animals was balanced in each cohort in each experiment. Ages of mice used in each experiment are indicated in the figures. Lung tumors were initiated by intratracheal intubation and delivery of 60 μl of lentiviral vectors in PBS to isoflurane-anesthetized mice. Lentiviral titers and durations of tumor growth are indicated in the figures.

Lungs were weighed at the time of collection and cancer cell weight (tumor-bearing lung weight minus normal lung weight) was calculated as a metric of overall tumor burden. Tumor burden was also assessed via fluorescence imaging and quantified using ImageJ. Lung lobes were frozen for DNA extraction and Tuba-seq analysis, fixed for histology, and/or dissociated into a single cell suspension for molecular analyses. Tissues were fixed in 4% formalin for 24 hours, stored in 70% ethanol, and paraffin-embedded. Hematoxylin and Eosin staining was performed on 4 µm thick sections by Histo-Tec Laboratory, Inc. using standard protocols.

Young and old mice used in analyzing the impact of age on the efficiency of lentiviral transduction of lung epithelial cells (**Fig. 1e** and **Extended Data Fig. 2**) were *C57BL/6* mice obtained from the National Institute of Aging’s aged rodent colony. These mice were housed in the Stanford SIM1 barrier facility for at least two months prior to experimentation.

### Transduction Efficiency Analysis

To assess the impact of age on lentiviral transduction efficiency of lung epithelial cells, young and old mice were intratracheally transduced with 1.0 x10^6^ ifu of a *GFP*-expressing lentiviral vector or with an equivalent volume of PBS (untransduced controls). To allow time for expression of GFP in transduced cells, lungs were collected 7-8 days after transduction. Lungs were inflated with digestion media containing collagenase IV, dispase, and trypsin and dissociated at 37°C for 30 minutes. Cells were stained with DAPI and antibodies against lineage markers CD45 (30-F11), CD31 (390), F4/80 (BM8), Ter119 and epithelial marker EpCAM (all from BioLegend). Transduction efficiency was measured by FACS for each sample via quantification of the percentage of live epithelial cells (EpCam^pos^Lineage^neg^DAPI^neg^) in each sample that were GFP-positive. Untransduced control samples were used to define an appropriate gating strategy.

### Tuba-seq library generation and sequencing

For Tuba-seq library generation, genomic DNA was isolated from whole tumor-bearing lung tissue from each mouse as previously described^7^. Three benchmark control ‘spike-in’ cell lines (10^5^ cells each) were added to each lung sample before lysis to enable the calculation of the absolute number of neoplastic cells in each tumor from the number of sgID-BC reads^1^. Following homogenization and overnight protease K digestion, genomic DNA was extracted from the lung lysates using phenol–chloroform and ethanol precipitation. Libraries were prepared by amplifying the sgID-BC region from 32μg of genomic DNA per mouse using unique dual-indexed primers and the Q5 Ultra II 2x Master Mix (New England Biolabs, M0544X). The PCR products were purified with Agencourt AMPure XP beads (Beckman Coulter, A63881). The concentration and quality of the purified libraries were determined using the Tapestation (Agilent Technologies).

Unequal sequencing depth can be a source of noise in characterizing the distribution of tumor sizes across samples, and a potential confounding factor if systematically different across experimental groups. To ensure that samples were sequenced evenly, we performed two sequential rounds of paired-end sequencing for each set of Tuba-seq samples. Sequencing depths were ascertained using the read counts of the spike-in cell lines in the first round of sequencing and were then used to adjust sample pooling for the second round to ensure equal overall sequencing depth. For both experiments using Lenti-*sgRNA^Aging^*/*Cre*, samples were divided into two sub-pools and sequenced on the Illumina MiSeq Nano, re-pooled, and then sequenced a second time on the Illumina NextSeq 500 platform (read lengths 2 x 150 bp, through Admera Health). To minimize the influence of batch effects associated with each sequencing lane, sub-pools were balanced for the sex and age of mice. For the individual gene validation experiments, samples were sequenced on a partial lane of Illumina NovaSeq 6000, re-pooled, and then sequenced on a second partial lane of NovaSeq 6000 (2 x 150bp, through Novogene) or were pooled on the basis of lung weight (a proxy for tumor burden). Three of the samples transduced with Lenti-*sgRNA^Aging^*/*Cre* had extremely aberrant sequencing depths and were removed from the analysis due to assumed failures in library preparation (an old *Kras^LSL-G12D/+^*; *H11^LSL-Cas9^* mouse, an old *Kras^LSL-G12D/+^*; *p53^flox/flox^*;*H11^LSL-Cas9^*mouse, and a *Kras^LSL-G12D/+^* mouse).

### Analysis of sgID-BC sequencing data

Sequencing of Tuba-seq libraries produces reads that are expected to contain an 8-nucleotide sgID followed by a 30-nucleotide BC of the form GCNNNNNTANNNNNGCNNNNNTANNNNNGC, where each of the 20 Ns represents a random nucleotide (Rogers et al). Each sgID has a one-to-one correspondence with an sgRNA in the viral pool (see **Supplemental Table 1**); thus, the sgID sequence identifies the gene targeted in each tumor. Note that all sgID sequences in a viral pool differ from each other by at least three nucleotides such that incorrect sgID assignment (and thus, inference of tumor genotype) due to PCR or sequencing error is extremely unlikely. The random 20-nucleotide BC tags all cells in a single clonal expansion. Note that the length of the BC ensures a high theoretical potential diversity, with the actual diversity of each Lenti-sgRNA/Cre vector dictated by the number of bacterial colonies pooled during the plasmid barcoding step.

FASTQ files were parsed using regular expressions to identify the sgID and BC in each read. To minimize the effects of sequencing error on BC identification, we required the forward and reverse reads to agree completely within the sgID-BC region^8^. We also performed an analysis to identify BCs that were likely to have arisen from genuine tumors due to PCR or sequencing errors. Given the low rate of sequencing error, we expect these ‘spurious tumors’ to have read counts that are far lower than the read counts of the genuine tumors from which they arise. To minimize the impact of these ‘spurious tumors’, we identified small ‘tumors’ with BCs that were highly similar to the BCs of larger tumors in the same sample. Specifically, if a pair of ‘tumors’ within a sample had BCs that were within a Hamming distance of two, and if one of the tumors had fewer than 5% as many reads as the other, then the reads associated with the smaller tumor were attributed to the larger tumor.

After these filtering steps, the read counts associated with each BC were converted to absolute neoplastic cell numbers by normalizing to the number of reads from the ‘spike-in’ cell lines added to each sample before lung lysis and DNA extraction.

### Removal of contaminating barcodes

sgID-BC sequences that are not from genuine tumors in an individual tumor-bearing lung sample can none-the-less be present in sequencing libraries for several reasons including intra-experiment sample-to-sample cross-contamination during library preparation, external contamination (*e.g*. from samples and libraries from other experiments), and library-to-library misassignment during sequencing. These sgID-BC sequences have the potential to be identified as small tumors (*i.e*. ‘spurious tumors’) and thereby reduce the precision of our analyses. Both external contamination and sample-to-sample contamination result in the identification of the same sgID-BC in multiple samples. However, some sgID-BCs are expected to recur across samples in the absence of contamination due to the finite diversity of the sgID-BC region in each lenti-sgRNA/Cre vector. To minimize the effects of contamination, we examined patterns of sgID-BC recurrence across samples and removed barcodes that occurred in a number of samples that would be highly unlikely to occur by chance given the BC diversity of each vector.

To estimate the BC diversity associated with each sgID, we assume that the probability of observing a BC in *i* mice is Poisson distributed: P(*k*=*i*; λ) = *λ*^k^ *e*^−*λ*^ */ k*!, where *λ* is the mean number of mice that barcodes appear in for a given lenti-sgRNA vector. To estimate *λ*_r_ for each sgID *r* in the dataset we note that *λ*_r_/(1 – *e^−λ^*^r^) = *μ*_non-zero_, where *μ*_non-zero_ = Σ^∞^P(*k=i*; *λ*_r_) is the mean number of occurrences of each BC that occurred once or more (a known quantity). Given the Poisson distribution defined for each sgID *r*, we calculated a lenti-sgRNA/Cre vector specific threshold N_r_ such that 99.9% of barcodes would be expected to appear in N_r_ samples or fewer and identified and removed all ‘tumors’ with sgID-BCs that occurred in a number of samples exceeding N_r_.

sgID-BCs can also recur across samples due to misassignment of reads during sample de-multiplexing. While misassignment of reads is expected to be extremely rare, it could result in sgID-BCs from genuine tumors with very large read counts in one sample (i.e., a very large, genuine tumor) appearing with low read counts in additional samples sequenced in the same lane. To guard against this possibility and ensure that very large tumors were not systematically discarded by the procedure described in the prior paragraph, we examined the distribution of reads across samples to identify sgID-BCs that appeared in >N_r_ samples but where >95% of sequencing reads were assigned to a single sample. These sgID-BCs were retained in the sample with the bulk of the sequencing reads and discarded in all other samples. All other barcodes appearing in > N_r_ samples were discarded from all samples.

### Removal of mice with aberrantly few barcodes

After processing sequencing data, the number of unique barcodes was tallied in each sample. Barcode number across mice was highly variable for both young and old mice, which likely reflects variability in the success of lentiviral transduction due to technical reasons. Samples with extremely low barcode number (fewer than 1,000 barcodes, compared to an average of 87,868 barcodes detected in mice transduced with Lenti-*sgRNA^Aging^*/*Cre*) were deemed to have not been successfully transduced and were removed from the analysis. On this basis, we removed one young and one old sample from the cohort of *Kras^LSL-G12D/+^*;*H11^LSL-Cas9^*transduced with Lenti-*sgRNA^Aging^*/*Cre* and two young samples from the cohort of transduced with *Kras^LSL-G12D/+^*;*p53^flox/flox^*;*H11^LSL-Cas9^*mice transduced with Lenti-*sgRNA^Aging^*/*Cre*.

### Filtering of vectors with insufficient titer from analysis

Although Lenti-sgRNA/Cre vectors were titered before pooling to aim for the desired representation in the Lenti-*sgRNA^Aging^*/*Cre* viral pool, the distribution of vectors in the viral pool was uneven. Examination of the number of tumors associated with each sgRNA in the *K;H11^LSL-Cas9^* mice revealed that a small number of vectors had titers roughly an order of magnitude lower than the median titer (<3,000 unique barcodes relative to the median of 46,793). Vectors with fewer than 3,000 unique barcodes identified across the young and old *K;H11^LSL-Cas9^* mice were deemed to be insufficiently represented and were filtered out prior to analysis. Critically, all filtered vectors were similarly poorly represented in the *Kras^LSL-G12D/+^* control mice (<750 unique barcodes per vector, relative to the median of 9,695), indicating that the low barcode counts in the experimental cohorts were due to uneven pooling rather than the impacts of gene targeting.

### Adaptive sampling of tumors for statistical comparison of tumor genotypes

In comparing the effects of tumor suppressor inactivation in young and old mice, we sought to compare equivalent portions of the tumor size distributions (*i.e*., the same number of tumors per infectious unit of virus delivered) for each sgRNA in the young and old cohort. To ensure this, we scaled the number of tumors analyzed for each sgRNA *i* in each cohort *j* to account for differences in viral titer and the number of mice transduced in each cohort, and then analyzed the largest *N_i_*_,*j*_ tumors per Lenti-sgRNA/Cre vector.

This scaling procedure requires selecting a benchmark sgRNA and a benchmark cohort, and then selecting a defined number of tumors with that benchmark sgRNA from that cohort. For each analysis, we used *sgNT1* in the young cohort as the benchmark dataset. Note that this choice was arbitrary, and that identical results could be achieved by basing the sampling calculations on any sgRNA in either the young or old cohort. The number of tumors sampled for each other sgRNA *i* in each cohort *j* (*N_i_*_,*j*_) is then adjusted to take into account the proportions of sgRNAs in the viral pool and differences in the overall viral titer delivered to the young and old cohorts:

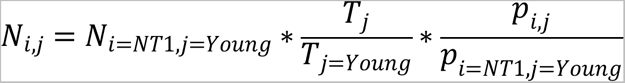

where *T_j_* denotes the total viral titer delivered to cohort *j* and *p_i_*_,*j*_ denotes the proportion of the viral pool in delivered to cohort *j* that is allocated to sgRNA *i*. *T_j_* is a known quantity determined during viral preparation which represents the sum of the viral titers delivered to each mouse in cohort *j*. *P_i_*_,*j*_ values were calculated using Tuba-seq data generated by transducing control *Kras^LSL-G12D/+^* mice. As these mice lack Cas9 expression all Lenti-sgRNA/Cre vectors are functionally inert, and the observed tumor number associated with each sgRNA reflects the make-up of the viral pool. To account for variation in tumor number across mice the sample of *N_i_*_,*j*_ tumors was distributed across all mice in the cohort in proportion to the total number of tumors in each mouse.

### Selection of number of tumors to sample in benchmark dataset (**N_i=NT1,j=Young_)**

The parameter *N_i_*_=*NT*1,*j*=*Young*_ defines the portion of the tumor size distribution that we use in assessing the impact of age on tumorigenesis and tumor suppressor function. In selecting *N_i_*_=*NT*1,*j*=*Young*_ for each analysis we sought to include as much data as possible while maintaining high data quality by excluding tumors with barcodes supported by very few reads. We reasoned that restricting our analysis to tumors supported by an average of >5 reads in each dataset would minimize noise associated with the detection and measurement of very small tumors while maintaining high statistical power to detect the impacts of aging on tumor suppressive effects. We therefore selected an *N_i_*_=*NT*1,*j*=*Young*_ for each analysis that corresponded to the number of sg*NT1* tumors supported by at least 5 reads in the young cohort (rounded to the nearest 100). These values of *N_i_*_=*NT*1,*j*=*Young*_ are shown in the table below. To ensure that our findings were robust to variation in the chosen values of *N_i_*_=*NT*1,*j*=*Young*_, we performed sensitivity analyses by increasing/decreasing *N_i_*_=*NT*1,*j*=*Young*_ by 10, 25, and 50% and found similar results (**Extended Data Fig. 6**).

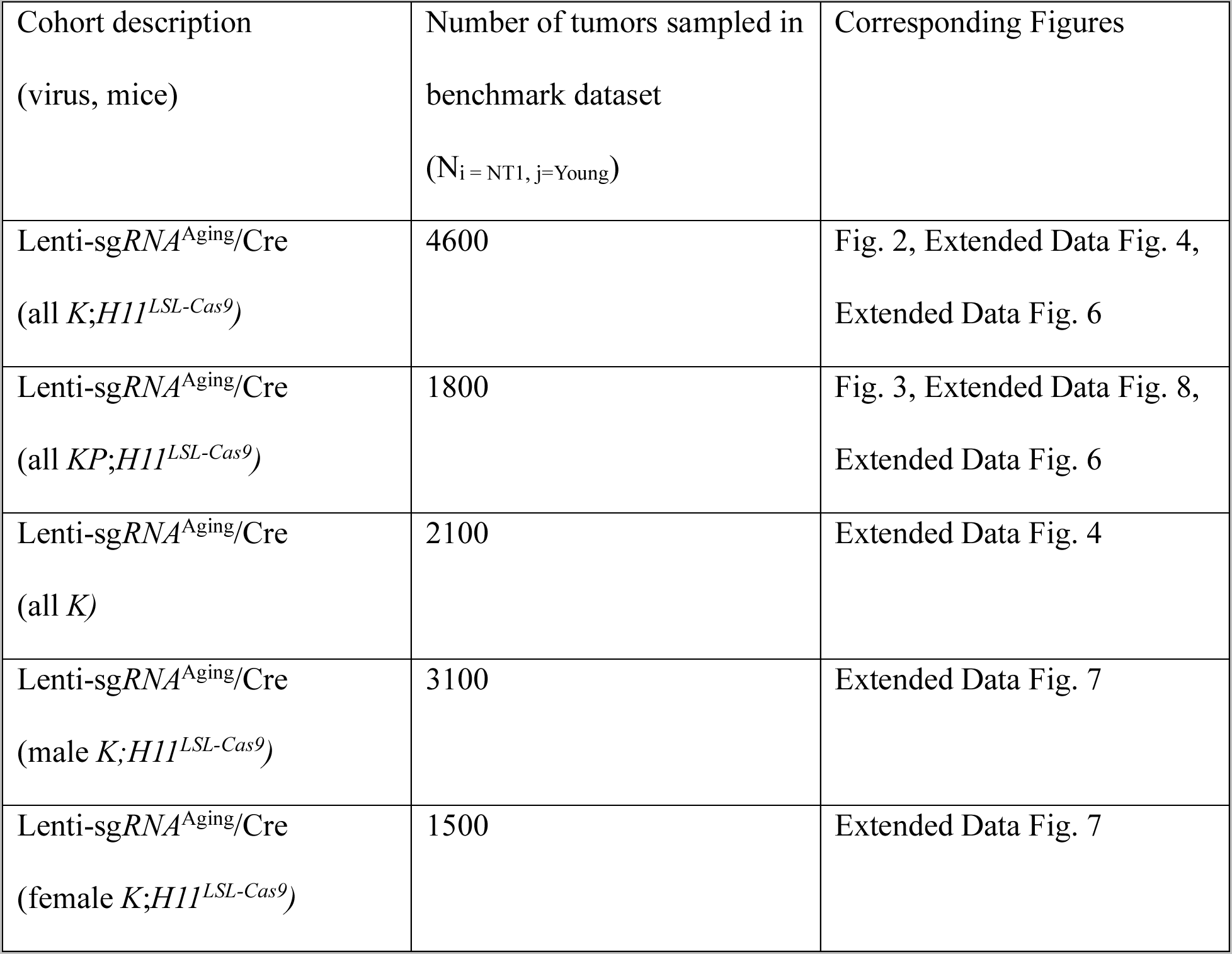
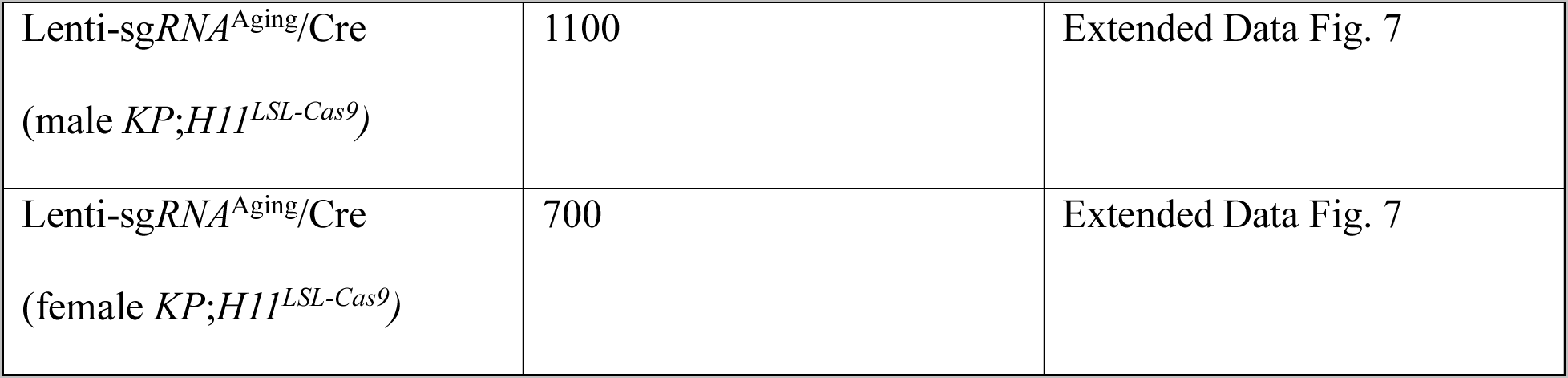

### Summary statistics for impact of gene inactivation on tumorigenesis

After performing the adaptive sampling described above, we assessed the extent to which each targeted gene (*X)* affects tumor initiation and growth, by comparing the distribution of tumor sizes produced by vectors targeting that gene (sg*X* tumors) to the distribution produced by our negative control vectors (sg*Inert* tumors). We characterized these distributions using the maximum-likelihood estimate of mean tumor size assuming a log-normal distribution (LN mean or ASM). Previous work found that this statistic represents the best parametric summary of tumor growth based on the maximum likelihood quality of fit of various common parametric distributions^1^. In addition, we calculated the sizes of tumors at various percentiles of the distribution (50^th^, 60^th^, 70^th^, 80^th^, 90^th^) for each sgRNA as complementary nonparametric summary statistics.

Note that while these statistics are functions of the tumor size distribution, they are also sensitive to the effects of tumor suppressor gene inactivation on tumor initiation due to our use of adaptive sampling. For example, a gene which increases tumor burden when inactivated solely by increasing the number of tumors (i.e., without shifting the tumor size distribution) would still increase the LN mean of the adaptively sampled datasets and would therefore still be detectable as a tumor suppressor using these metrics. These statistics therefore integrate the impact of each targeted gene on the initiation and growth of tumors.

To quantify the extent to which each gene suppressed or promoted tumorigenesis, we normalized statistics calculated on tumors of each genotype to the corresponding sg*Inert* statistic. The resulting ratios reflect the fitness advantage (or disadvantage) associated with each tumor genotype relative to the initiation and growth of *sgInert* tumors.

The adaptively sampled relative LN mean size for tumors of genotype X was calculated as:

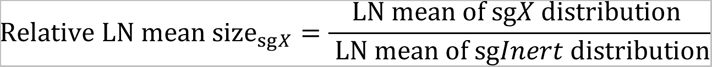

Likewise, the adaptively sampled relative i^th^ percentile for tumors of genotype X was calculated as:

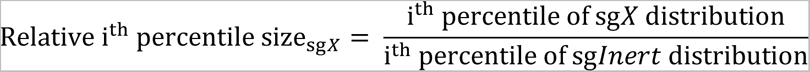

where the sg*X* and sg*Inert* distributions were adaptively sampled as previously described and the sg*Inert* statistic is the median value across all inert vectors.

### Aggregating information across sgRNAs in calculation of tumor growth metrics

Adaptive sampling and calculation of summary statistics was performed per sgRNA. To calculate statistics at the gene level, we aggregated information across all sgRNAs targeting the same gene. Specifically, the gene-level statistics reported are the weighted average of the per-sgRNA statistics, where the contribution of each sgRNA was proportional to the number of adaptively sampled tumors with that sgRNA.

### Calculation of confidence intervals and P-values for impacts of gene inactivation on tumorigenesis

Confidence intervals and *P*-values were calculated using bootstrap resampling to estimate the sampling distribution of each statistic. To account for both mouse-to-mouse variability and variability in tumor size and number within mice, we used a two-step, nested bootstrap approach where we first resampled mice, and then resampled tumors within each mouse in the pseudo-dataset. 10,000 bootstrap samples were drawn for all reported P-values. 95% confidence intervals were calculated using the 2.5^th^ and 97.5^th^ percentiles of the bootstrapped statistics. Because we calculate metrics of tumor growth/burden that are normalized to the same metrics in sg*Inert* tumors, under the null model where genotype does not affect tumor growth the test statistic is equal to 1. Two-sided p-values were thus calculated as followed:

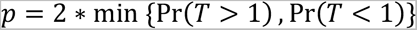

Where T is the test statistic and Pr(T>1) and Pr(T<1) were calculated empirically as the proportion of bootstrapped statistics that were more extreme than the baseline of 1. To account for multiple hypothesis testing, p-values were FDR-adjusted using the Benjamini-Hochberg procedure as implemented in the Python package stats models or in R Stats package.

### Calculation of p-values for differential effects of tumor suppressor inactivation with age

To compare the impacts of inactivating a given tumor suppressor gene in young and old mice, we defined a test statistic T equal to the difference between the effect in young and old. We produced a bootstrapped distribution of T by resampling mice and tumors within each cohort (as described above) and repeatedly calculating T. Under the null hypothesis, the impact of inactivating a given tumor suppressor gene is the same in young and old and T is therefore equal to 0. Two-sided p-values were thus calculated as followed:

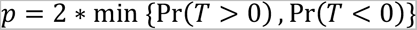

Where Pr(T>0) and Pr(T<0) were calculated empirically as the proportion of bootstrapped statistics that were more extreme than the baseline of 0. To account for multiple hypothesis testing, p-values were FDR-adjusted using the Benjamini-Hochberg procedure as implemented in the Python package stats models or in R Stats package.

### Preparation of bulk tumor-bearing lungs for scRNA-sequencing

Whole tumor-bearing lungs were finely minced immediately after harvesting and enzymatically dissociated using collagenase IV, dispase, and trypsin at 37°C for 30 minutes as previously described^9^. Bulk cell suspensions were viably frozen in Bambanker cryopreservation media (Wako Chemicals, 302-14681) at −80°C before transfer to liquid nitrogen. Cells were thawed and stained with antibodies against CD45 (30-F11), CD31 (390), F4/80 (BM8), and Ter119 (all from BioLegend) to identify hematopoietic and endothelial cells. Each sample was sorted for viability (DAPI-negative) and divided into two compartments: (1) “cancer cell” samples containing Tomato^pos^, CD45^neg^, CD3^neg^, F4/80 ^neg^ and Ter119 ^neg^ cells which are presumed to be cancer cells, and (2) “bulk” samples containing all other viable cells. FACSAria™ sorters (BD Biosciences) were used for all cell sorting. Samples were fixed immediately after sorting using Parse Biosciences Cell Fixation kits (v2, May 2023) and stored at −20°C. To maximize recovery for cancer cell samples where we had limited input material, we used a modified version of the Parse fixation protocol. Specifically, all volumes were scaled down to ¼ of the recommended amount so that the entirety of the final suspension of fixed cells could be loaded into the plate used for barcoding and library preparation.

### Single-cell RNA sequencing

Fixed cell suspensions were thawed and visually inspected under the microscope for the presence of debris, and one bulk sample was excluded on this basis (from mouse MT1906). The remaining 31 samples were processed for multiplexed single-cell RNA-sequencing using the Parse Biosciences WT Mega v2 kit according to the manufacturer’s instructions. This kit uses a combinatorial barcoding approach, allowed multiplexing of all samples into 16 sublibraries, each containing a mixture of cells from all samples. Samples for this experiment were loaded into the kit and processed alongside and sequenced with samples from another experiment. Sublibraries were quantified by Qubit fluorometer, pooled to equalize the amount of DNA per cell, and sequenced on an Illumina NovaSeq S4 flow cell to an average depth of 25,901 reads per cell.

### scRNA sequencing data processing and quality control

The processing pipeline provided by Parse Biosciences (split-pipe v1.1.1) was used with default settings to align sequencing reads to the GRCm38 mouse reference genome. Each of the 16 sublibraries was first processed individually using the command split-pipe –mode all, and the output of the 16 sublibraries was combined using split-pipe –mode combine.

Cell barcodes were filtered to include only cells meeting the following criteria: (1) expressing between 200 and 10,000 genes, (2) having between 500 and 100,000 read counts, and (3) having less than 10% of reads map to mitochondrial genes. Scrublet was used to simulate doublets and assign doublet scores to cells based on a k-nearest-neighbor classifier^10^. Cells with doublet scores >0.12 were removed from the dataset on the basis of visual inspection of the doublet score density plot for simulated doublets and inspection of UMAP embeddings. Read counts were normalized per cell using the scanpy function ‘normalize_total’ with default parameters and scaled for variance stabilization using a shifted logarithm transformation (log X+1, where X is the normalized counts).

### Dimensionality reduction, clustering, and visualization of single cell profiles

After filtering and normalization, feature selection was performed on the logarithmized data to identify transcriptionally overdispersed genes. Specifically, the scanpy function ‘highly_variable_genes’ was used to bin genes based on their mean expression and then annotate genes with high levels of dispersion relative to genes in the same bin. UMAP embedding was then performed to visualize single cell profiles in two-dimensional space^11^. Leiden clustering was performed at various resolutions to aid in cell type assignment (see below)^12^.

### Cell type assignment

Cell types were assigned after quality control and doublet removal on the basis of marker gene expression using Semi-supervised Category Identification and Assignment (SCINA^13^). Cell type identities and corresponding marker genes were taken from the LungMAP Consortium’s Mouse cell reference, accessed through the LGEA Web Portal (https://research.cchmc.org/pbge/lunggens/CellRef/LungMapCellRef.html)^14,15^. For each cell type, all genes listed as positive signatures in LungMAP were used as marker genes. SCINA was run using default model parameters.

SCINA assigns cell types to each cell individually. To assign types to cells that were not successfully annotated (i.e. annotated with type “unknown”) and ensure homogenous cell type assignments within clusters, we refined this initial clustering through two rounds of cluster-based annotation. In the first round, a coarse-grained clustering was performed to separate cells by broad lineage identity (epithelial, mesenchymal, endothelial, myeloid and lymphoid).

Specifically, cells were clustered using the Leiden algorithm implemented in scanpy with a resolution parameter of 0.5^12^. Each cluster was assigned a lineage identity based on the majority lineage identity of its constituent cells (where the mapping of cell type to lineage identity was specified by LungMAP cell reference). The dataset was then subdivided into epithelial cells, mesenchymal cells, endothelial cells, myeloid cells and lymphoid cells. A finer-grained clustering with a resolution parameter of 1.0 was then performed within each these broad cell types. Final cell types were then assigned on a per-cluster basis based on majority cell type of the constituent cells in each cluster.

After automatic annotation, cell type annotations were manually reviewed. Clusters with membership equally spread across two or more cell types were annotated with a more generic cell type label encompassing the component cell types. Specifically, clusters containing mixtures of T regulatory, CD4+ T cells, CD8+ T cells and NK cells were annotated as T lymphocytes/NK cells; clusters containing mixtures of alveolar fibroblast 1 and 2 cells were annotated as alveolar fibroblasts; and mixtures of endothelial cells and capillary cells were annotated as endothelial cells. In the rare instances where this was not possible, clusters were annotated as “unknown” and removed from downstream analysis.

### Removal of transduced non-cancer cells

Although lentivirus primarily has tropism for alveolar epithelial cells, we expect rare transduction of other cell types^16^. Although these events are not expected to give rise to tumors, they could impact the molecular state of the transduced cells. To minimize these effects we removed all non-epithelial cells that either expressed *tdTomato* transcripts or were sorted into cancer cell samples during our sample preparation for single-cell RNA-sequencing (and therefore have a higher likelihood of expressed Tomato at the time of sorting).

### Definition of cancer cell population

Lung adenocarcinomas arise from cells in the alveolar epithelium^17,18^. Consistent with this, *Tomato* expression in our dataset mapped primarily to AT1-, AT2- and AT/AT1-like cells. It is, however, likely that a portion of these alveolar cells were not transduced. While in theory *Tomato* expression serves as a marker of transduction, expression of the transgene was insufficiently high to serve as a reliable marker of cancer cell identity (i.e., exclusion on the basis of failure to express *Tomato* would discard a large number of likely cancer cells) (**Extended Data Fig. 10**). We reasoned that AT1, AT2, and AT1/AT2-like cells that did not express *Tomato* and were sorted into “bulk” samples were especially likely to be true-negatives and were enriched for untransduced alveolar epithelial cells. We therefore removed these cells (which accounted for ~4% of all alveolar epithelial cells) prior to downstream analyses and considered the remaining AT1-, AT2- and AT1/AT2-like cells to be our cancer cell population.

### Differential gene expression analysis

Cell type-resolved differential gene expression analysis was performed using a pseudobulk approach to avoid potential issues stemming from psuedoreplication^19,20^. Psuedobulk aggregation was performed by summing unnormalized read counts per gene for each cell type in each sample using the decoupler function get_pseudobulk^21^. Pseudobulked samples corresponding to fewer than 10 cells or 1000 counts were removed, and only cell types with at least two psuedobulked samples for each genotype-age category were analyzed.

Differential gene expression analysis was performed using DESeq2^22^ with unnormalized psuedobulked counts as input. Count tables were filtered separately for each cell type to include only genes with at least 10 counts across at least 3 samples. We performed two sets of analyses for each cell type: the first tested the effect of age on gene expression for each cell type separately within the sg*Inert* and sg*Pten* samples (Expression ~ Age); the second tested the effect of *Pten* inactivation on gene expression separately within the young and old samples (Expression ~ Pten^−/-^). For all analyses we used the standard DESeq2 workflow with log fold change shrinkage and reported differentially expressed genes as those with Benjamini Hochberg-corrected Wald test p < 0.05.

### Signatures of normal (non-cancer) aging

To assess the extent to which the transcriptional changes in our dataset aligned with pre-existing data on normal (non-cancer) aging, we identified three pre-existing aging signatures: (1) genes upregulated with age in AT2 cells in a single cell transcriptomic analysis of mouse lung (“Angelidis”)^23^, (2) genes upregulated with age in AT2 cells in the Tabula Muris Senis (“Tabula Muris”)^24^, and (3) genes that are upregulated with age in >80% of cell types in a pan-cell type re-analysis of the Tabula Muris Senis dataset (“Global Aging”)^25^. We assessed agreement of these signatures with the transcriptional changes in our dataset through gene set enrichment analyses and per-cell gene set activation scoring (see below).

### Gene set enrichment analysis and gene set activation scoring

Gene set enrichment analysis (GSEA) was performed per cell type using GSEA v4.3.2 with default parameters^26,27^. Genes were ranked by −log10(Wald p-value) *sign(log fold change) from DESeq2, and compared with gene sets corresponding to signatures of normal aging (see above)

To complement our GSEA analyses, which are based on the results of differential gene expression analysis using pseudobulked samples, we also performed per-cell scoring of aging gene sets. Specifically, we used the score_genes method implemented in scanpy, which quantifies the extent to which a gene set is active in each cell by subtracting the average expression of a reference set of genes randomly sampled across binned expression levels from the average expression of the gene set.

### Pathway activation analysis

PROGENy was used to estimate changes in signaling pathway activity with age and *Pten* inactivation based on changes in gene expression associated with these perturbations. This approach models signaling pathways as sets of target genes with interaction weights corresponding to the direction and extent of change in expression expected upon pathway activation (Schubert et al 2018). For each perturbation, we used the decoupler python package to fit a multivariate linear model predicting DESeq2 log2 fold changes in gene expression based on PROGENy pathway-gene weights^21^. Reported pathway activity scores are the t-values for the model coefficients for each pathway.

### Statistics and reproducibility

The statistical tests used for each analysis are described in detail in the sections above. All analyses of barcode sequencing data were performed in Python (3.6.4) and visualizations of data were performed in Python (3.6.4) and R (4.3.2). Sample sizes were determined on the basis of our previous experience conducing similar experiments and, in the case of barcode sequencing experiments, on the basis of previously published power analyses ^2^. Analyses of barcode sequencing data used nonparametric statistics; therefore, no assumptions about the distribution of data were made. Other metrics of tumorigenesis (e.g. lung weight, tumor burden, tumor number) were compared using Wilcoxon-rank sum tests. Two-way ANOVAs were used to analyze interactions between age and tumor genotype, and homogeneity of variance was confirmed by Levene’s test and normality was confirmed by the Shapiro-Wilk test.

## Data availability

All barcode sequencing datasets will be made publicly available through the NCBI’s Sequence Read Archive Database and accession numbers provided prior to publication of the manuscript. All single-cell RNA sequencing data will be made publicly available through the Gene Expression Omnibus and accession numbers provided prior to publication of the manuscript.

## Code availability

The code used for data analysis in this study will be made available on GitHub prior to publication of the manuscript.

## ACKNOWLEDGEMENTS

We thank the Stanford Shared FACS Facility for flow cytometry services, the Stanford Veterinary Animal Care Staff for expert animal care, and the NIA Aged rodent colony. We thank Anne Brunet, Joe Lipsick, Belal Yasin, and members of the Winslow and Petrov laboratories for helpful comments and experimental support. E.G.S was supported by a Tobacco-Related Disease Research Program (TRDRP) predoctoral fellowship (T33DT6556). H.C. was supported by a Tobacco-Related Disease Research Program (TRDRP) fellowship (28FT-0019). J.D.H was supported by an American Cancer Society postdoctoral fellowship (PF-21-112-01-MM) and a Tobacco-Related Disease Research Program (TRDRP) fellowship (T31FT1619). Y.J.T is supported by the Canadian Institute of Health Research postdoctoral fellowship (CIHR MFE 176568). S.K was supported by National Cancer Institute’s F99/K00 CA234962. This work was supported by R01-CA234349 (to D.A.P and M.M.W), R01-CA230025 (to M.M.W), U01-AG077922 (to M.M.W.), and in part by the Stanford Cancer institute support grant (NIH P30-CA124435).

## AUTHOR CONTRIBUTIONS

E.G.S, M.M.W and D.A.P conceived the project and designed the experiments. E.G.S led experimental data production with contributions from S.K, M.K.T, J.D.H, Y.J.T, L.A, M.W, C.R.D, H.C and R.T. E.G.S performed all data analysis. M.M.W and D.A.P oversaw the project. E.G.S, D.A.P and M.M.W wrote the manuscript with input from all authors.

## COMPETING INTERESTS

D.A.P and M.M.W are founders and hold equity in Guide Oncology.

---

**Extended Data Fig. 1.**
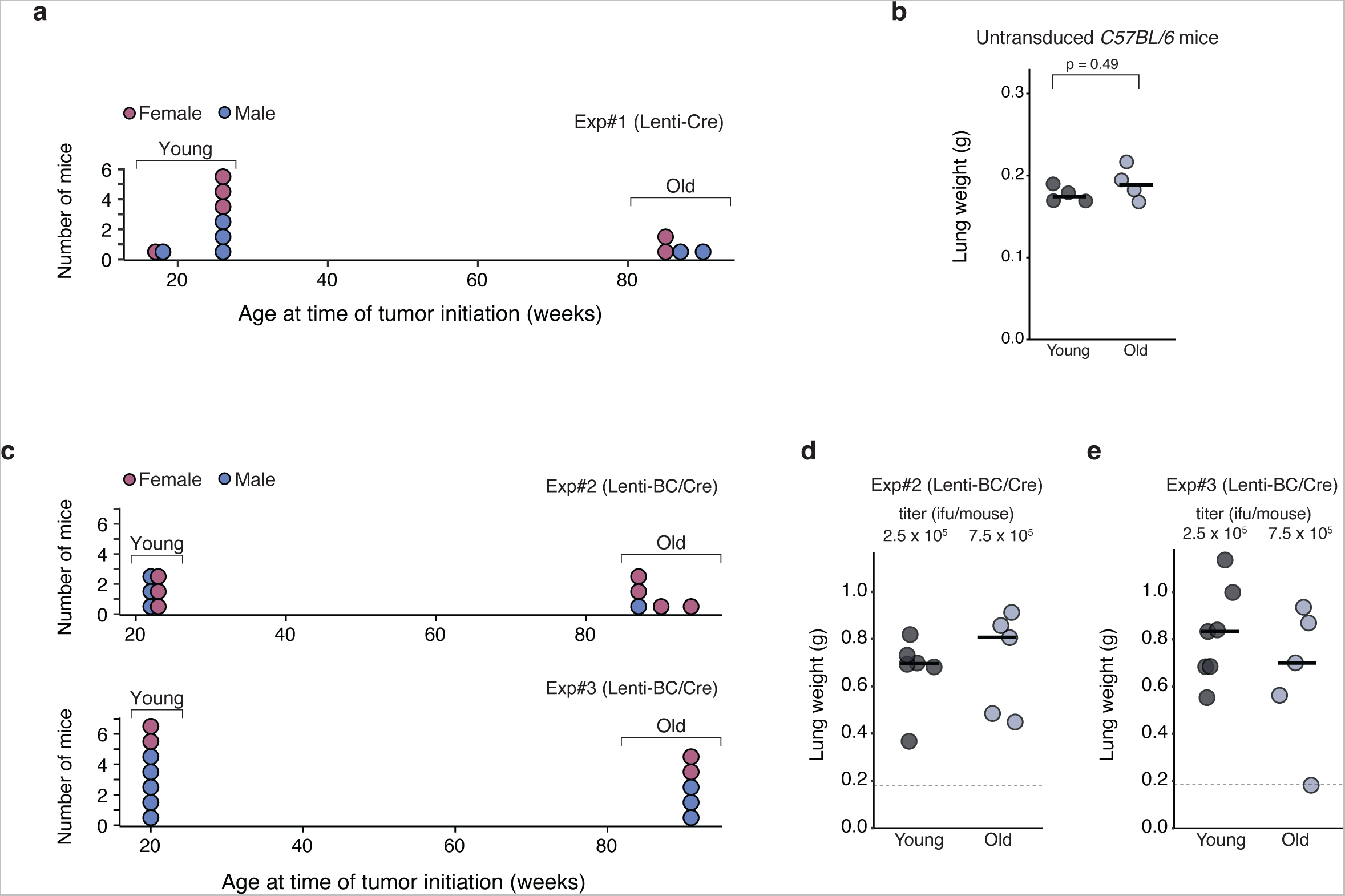
Additional data on generation of KRAS-driven tumors in young and old mice. **a.** Ages of young and old mice transduced with Lenti-Cre in Exp #1 (**Fig. 1a**). Each dot is a mouse; color denotes sex. **b.** Lung weights of untransduced young and old control mice matched to the ages of mice in Exp #1. Each dot is a mouse and the bars indicate the median values. **c.** Ages of young and old mice transduced with Lenti-BC/Cre in Exp #2 (top) and Exp #3 (bottom) (**Fig. 1f**). Each dot is a mouse; color denotes sex. **d,e**. Lung weights of mice transduced with Lenti-BC/Cre in Exp #2 (**d**) and Exp #3 (**e**). Each dot is a mouse and the bars indicate the median values. Median normal lung weight from (**b**) is indicated with dashed line.

**Extended Data Fig. 2.**
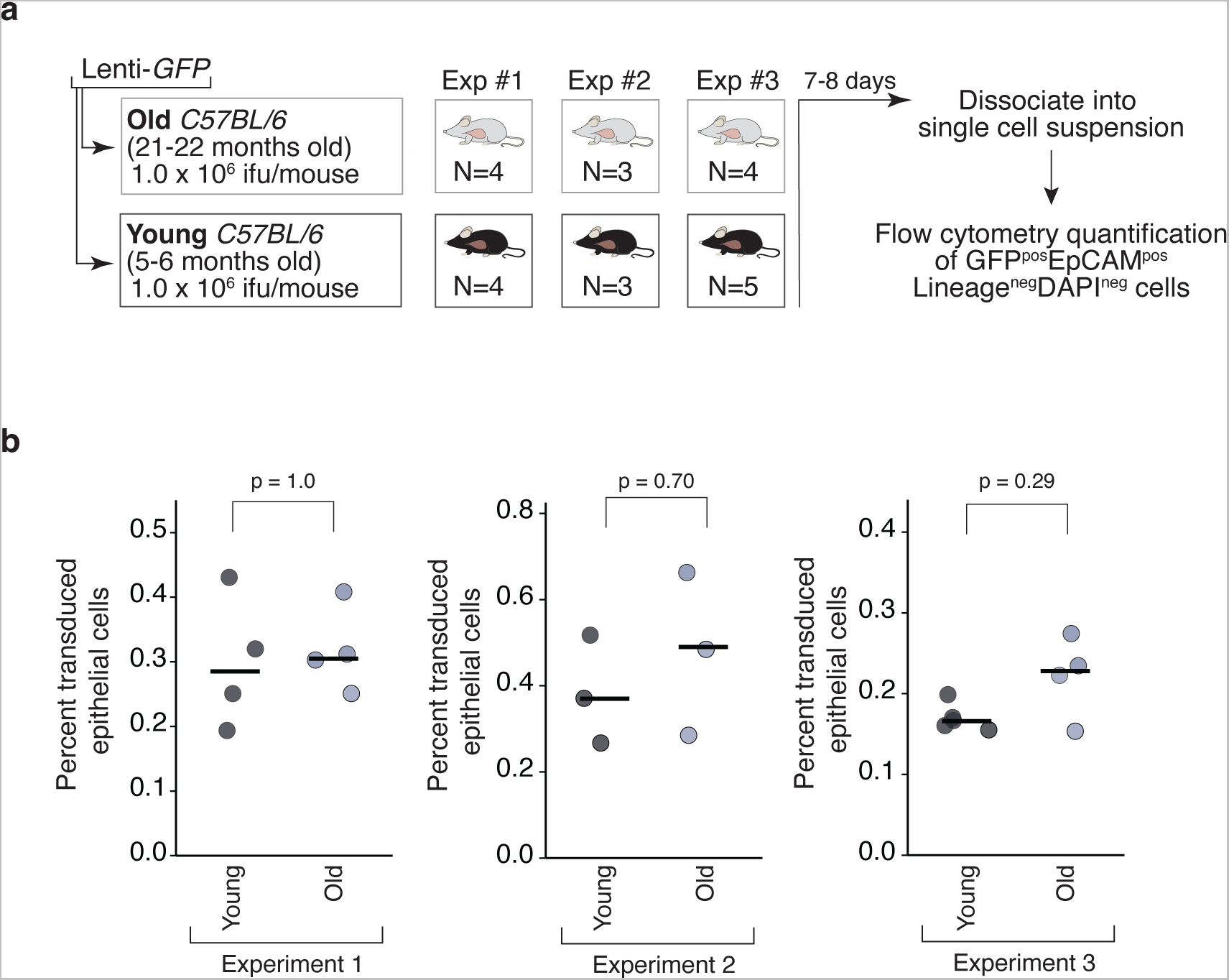
Lentiviral transduction efficiency of lung epithelial cells is not impacted by age. **a.** Transduction of young and old *C57BL/6* mice with Lenti-*GFP* to quantify lentiviral transduction efficiency. Lentiviral titer is indicated. Mice were analyzed 7-8 days after transduction. Lungs were dissociated and stained for EpCAM and lineage markers (CD31, CD45, and F4/80) to identify epithelial cells, as well as DAPI to exclude dead cells. Age and number of mice are indicated. **b.** Quantification of the percent of EpCAMposLineagenegDAPIneg cells that were GFPpos (transduced) in young and old mice across three replicate experiments. Each dot represents a mouse and the bars indicate the median values. P values, two-sided Wilcoxon rank sum tests.

**Extended Data Fig. 3.**
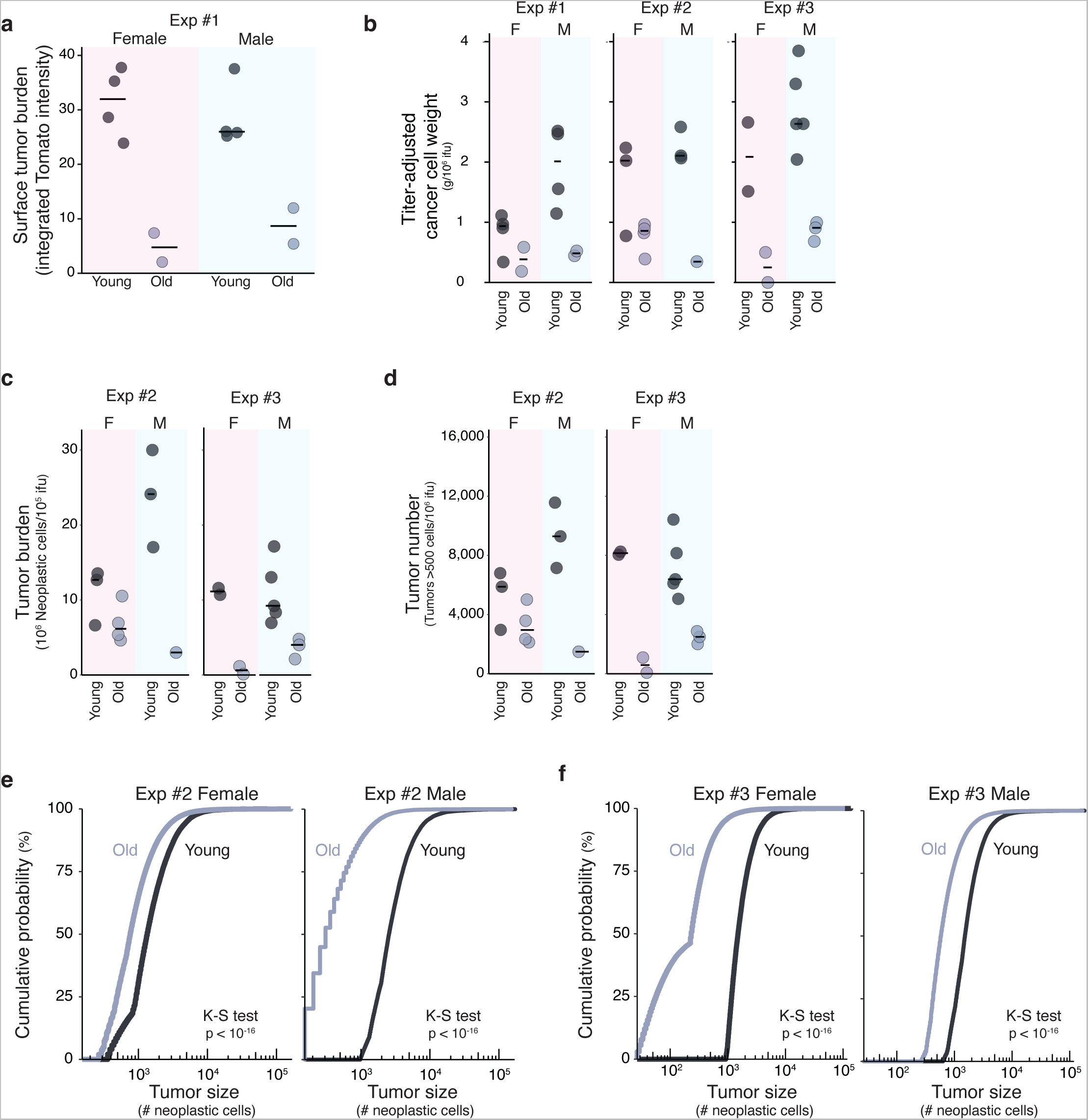
Aging represses KRAS-driven lung tumor initiation and growth in both males and females. **a.** Fluorescence-based quantification of tumor burden in young and old mice in Experiment #1 (transduced with Lenti-*Cre*) split by sex. Each dot is a mouse and the bars indicate median values. **b.** Estimated cancer cell weights of young and old mice in Experiments #1-3 (transduced with Lenti-*Cre* and Lenti-*BC/Cre*), normalized to the viral titer delivered to each mouse and split by sex. Each dot is a mouse and the bars indicate median values. **c,d**. Tumor burden (total number of neoplastic cells in clonal expansions > 500 cells quantified by Tuba-seq, **c**) and number of tumors (clonal expansions >500 cells, **d**) in young and old mice in Experiments #2 and #3 transduced with with Lenti-*BC/Cre* normalized to the viral titer delivered to each mouse and split by sex. Each dot is a mouse and the bars indicate median values. **e,f.** Empirical cumulative distribution functions of tumor sizes in young and old mice from Experiment #2 (**e**) and Experiment #3 (**f**) split by sex. To account for the 3-fold higher titer delivered to the old mice, this comparison includes the 10,000 largest tumors in each young sample and the 30,000 largest tumors from each old sample. Tumors in old mice are smaller than tumors in young mice irrespective of sex. K-S test: two-sided asymptotic Kolmogorov-Smirnov test.

**Extended Data Fig. 4.**
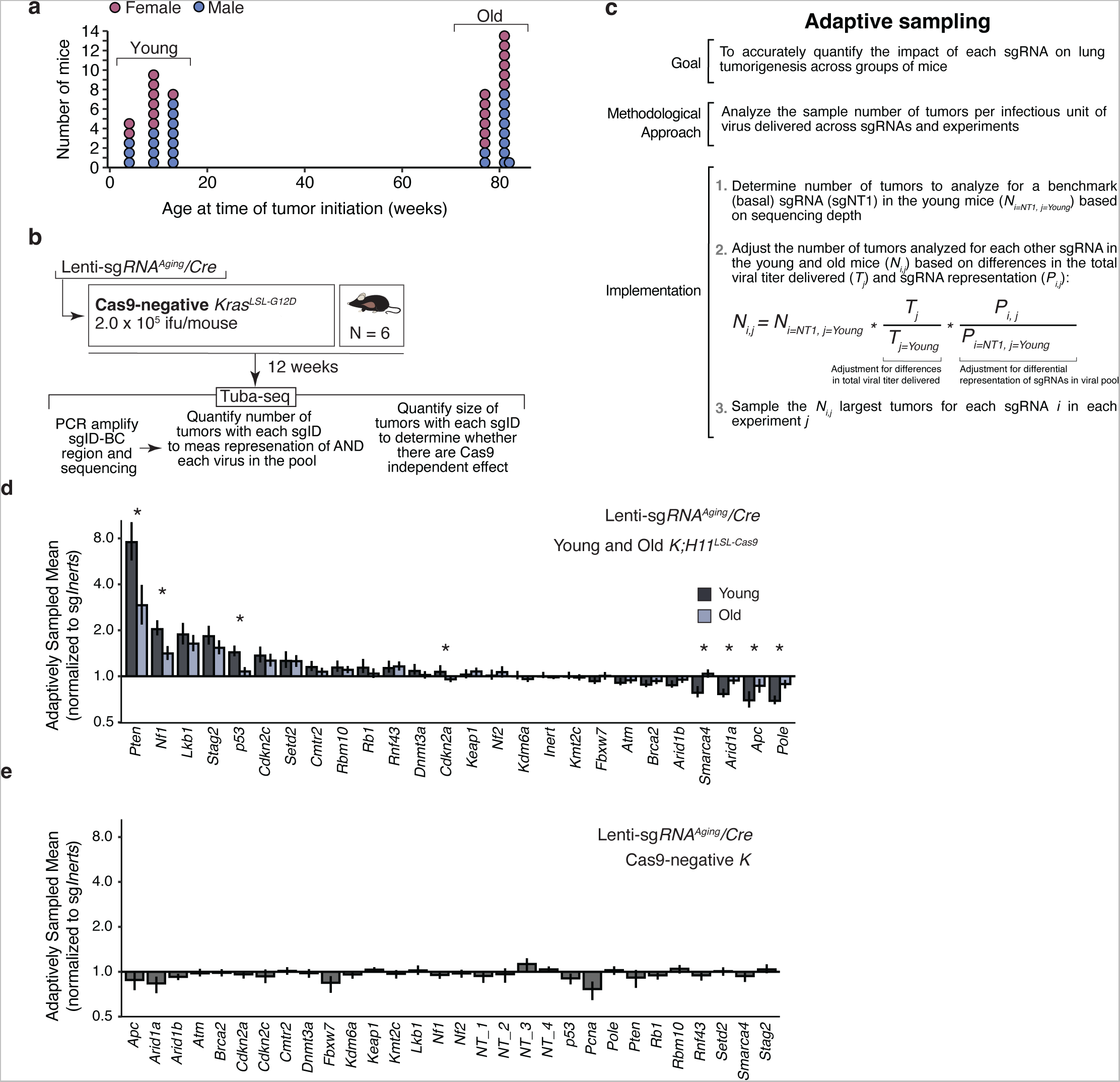
Aging impacts the effects of tumor suppressor gene inactivation in *K;H11LSL-Cas9 mice*. **a.** Ages of *K;H11LSL-Cas9* mice transduced with Lenti-*sgRNAAging*/Cre. Each dot is a mouse. Color denotes sex. **b.** Schematic of tumor initiation in *KrasLSL-G12D* (Cas9-negative) mice with the Lenti-sg*RNAAging/Cre* viral pool to determine representation of Lenti-sg*RNA/Cre* vectors. **c.** Explanation of the adaptive sampling method used in assessing the impact of Lenti-sg*RNA/Cre* vectors in young and old mice (see **Methods**). **d.** Adaptively sampled mean size (ASM) of each tumor genotype normalized to the ASM of sg*Inert* tumors in the young and old cohorts of *KrasLSL-G12D;H11LSL-Cas9* mice transduced with Lenti-sg*RNAAging*/Cre. ASM is a summary metric of tumor fitness that integrates the impact of inactivating each gene on tumor size and number. Line at x=1 indicates no effect on tumorigenesis. Stars denote a statistically significant difference (two-sided FDR-adjusted p-value < 0.05) between young and old. P-values and confidence intervals were calculated using nested bootstrap resampling. Genes are ordered by ASM in the young cohort. **e.** Adaptively sampled mean tumor sizes relative to sg*Inert* vectors for each sgRNA in the Lenti-sg*RNAAging/Cre* viral pool in Cas9-negative *KrasLSL-G12D* mice.

**Extended Data Fig. 5.**
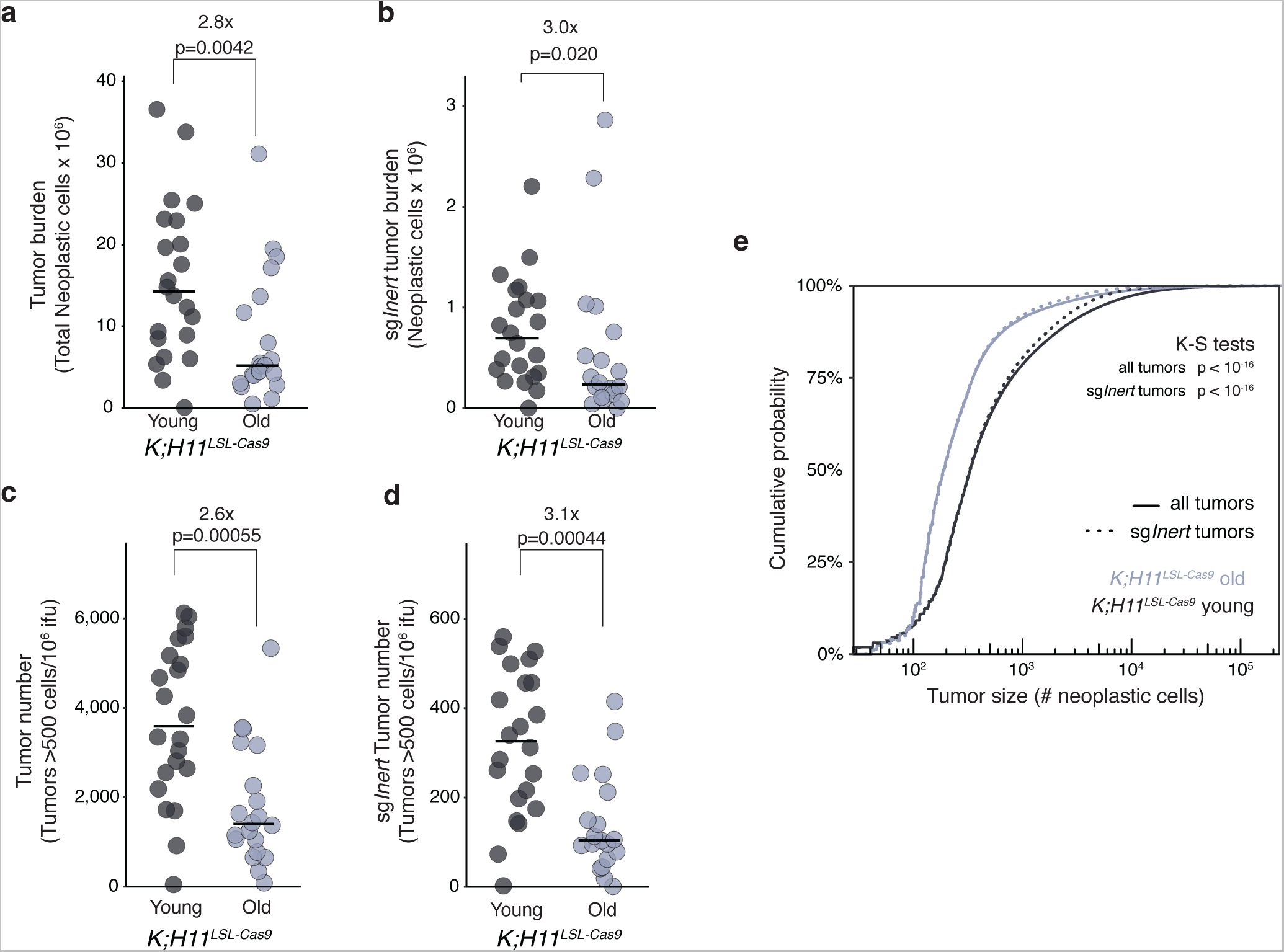
Pooled screen of tumor suppressor function recapitulates suppressive effect of aging on KRAS-driven lung tumorigenesis. **a-b.** Tumor burden (total number of neoplastic cells in clonal expansions > 500 cells) across all Lenti-sg*RNA/Cre* vectors (**a**) and across only sg*Inert* vectors (**b**) in young and old *K;H11LSL-Cas9* mice transduced with the Lenti-sg*RNAAging/Cre* pool. **c-d.** Number of tumors (clonal expansions > 500 cells) across all Lenti-sg*RNA/Cre* vectors (**e**) and across only sg*Inert* vectors (**f**) in young and old *K;H11LSL-Cas9* mice transduced with the Lenti-sg*RNAAging/Cre* pool.For **a**-**d**: Each dot is a mouse. Bars indicate median value within each age group. P-values were calculated using two-sided Wilcoxon rank sum tests. **e.** Cumulative distribution functions of tumor size across all Lenti-sg*RNA/Cre* vectors (”all tumors”) and across Lenti-sg*Inert/Cre* vectors (”sg*Inert* tumors”) in young and old *K;H11LSL-Cas9* mice transduced with the Lenti-sg*RNAAging/Cre* pool. Distribution for all tumors was constructed by sampling the 10,000 largest tumors per young and old mouse (irrespective of Lenti-sg*RNA/Cre* vector); distribution for *sgInert* tumors was constructed using the subset of the 10,000 largest tumors per mouse that contained Lenti-sg*Inert/Cre* vectors. K-S tests: two-sided asymptotic Kolmogorov-Smirnov tests comparing the overall distribution and the distribution of sg*Inert* tumor sizes between youn and old mice.

**Extended Data Fig. 6.**
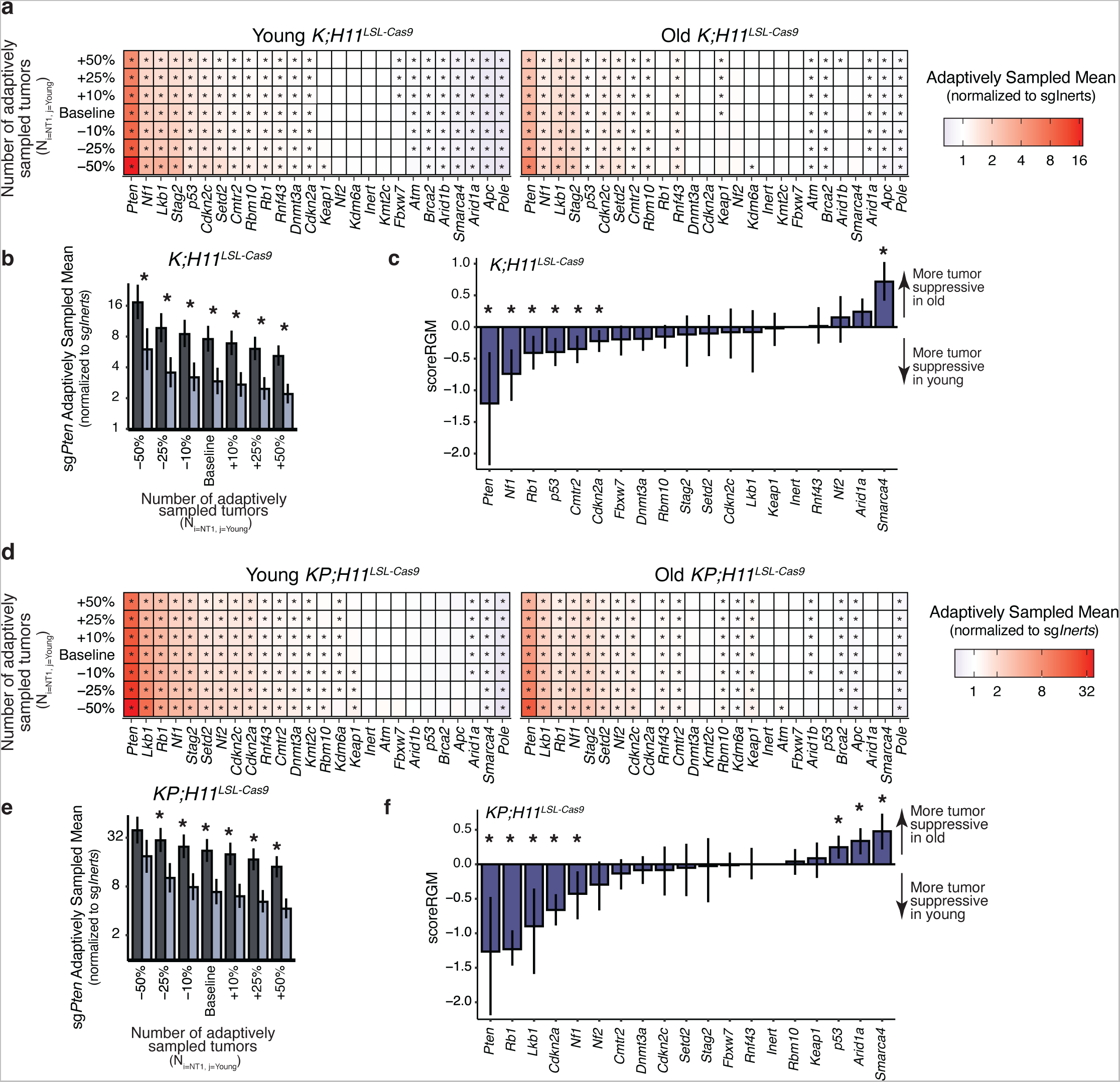
Differential effect of *Pten* inactivation with age is robust to variation in number of tumors adaptively sampled and statistics used to quantify impact of age on tumor suppressor function. **a.** Adaptively sampled mean (ASM) tumor sizes normalized to sg*Inert* for all tumor genotypes in young and old *K;H11LSL-Cas9* mice transduced with the Lenti-sgRNAAging/Cre pool, varying the number of tumors adaptively sampled. Stars denote a significant impact on ASM (two-sided FDR-corrected p<0.05). **b.** The impact of sg*Pten* on tumorigenesis is greater in young than old *K;H11LSL-Cas9* mice transduced with the Lenti-sg*RNAAging/Cre* pool across a range of adaptively sampled tumor numbers. Stars indicate a differential effect with age (FDR-adjusted two-sided p-value <0.05). For **a**-**b**: “Baseline” adaptively sampled tumor number corresponds to analyses shown in **Fig. 2**. Baseline adaptively sampled tumor numbers were calculated based on sequencing depth, viral titer, and the number of mice in each experiment (see **Methods**). **c.** Differential effects of tumor suppressor inactivation on tumor size with age as quantified by scoreRGM (Li *et al*.) in *K;H11LSL-Cas9* mice transduced with Lenti-sgRNAAging/Cre pool. Stars indicate differential effect with age (FDR-adjusted two-sided p-value <0.05). **d.** ASM tumor sizes normalized to sg*Inert* for all tumor genotypes in young and old *KP;H11LSL-Cas9* mice transduced with the Lenti-sgRNAAging/Cre pool, varying the number of tumors adaptively sampled. Stars denote a significant impact on ASM (two-sided FDR-corrected p<0.05). **e.** The impact of *sgPten* on tumorigenesis is greater in young than old *KP;H11LSL-Cas9* mice transduced with the Lenti-sgRNAAging/Cre pool across range of adaptively sampled tumor numbers. Stars indicate a differential effect with age (FDR-adjusted two-sided p-value <0.05). For **d**-**e**: “Baseline” adaptively sampled tumor number corresponds to analyses shown in **Fig. 3**. Baseline adaptively sampled tumor numbers were calculated based on sequencing depth, viral titer, and the number of mice in each experiment (see **Methods**). **f.** Differential effects of tumor suppressor inactivation on tumor size with age as quantified by scoreRGM (Li *et al*.) in *KP;H11LSL-Cas9* mice transduced with Lenti-sgRNAAging/Cre pool. Stars indicate differential effect with age (FDR-adjusted two-sided p-value <0.05) **All panels**: Confidence intervals and P-values calculated by nested bootstrap resampling.

**Extended Data Fig. 7.**
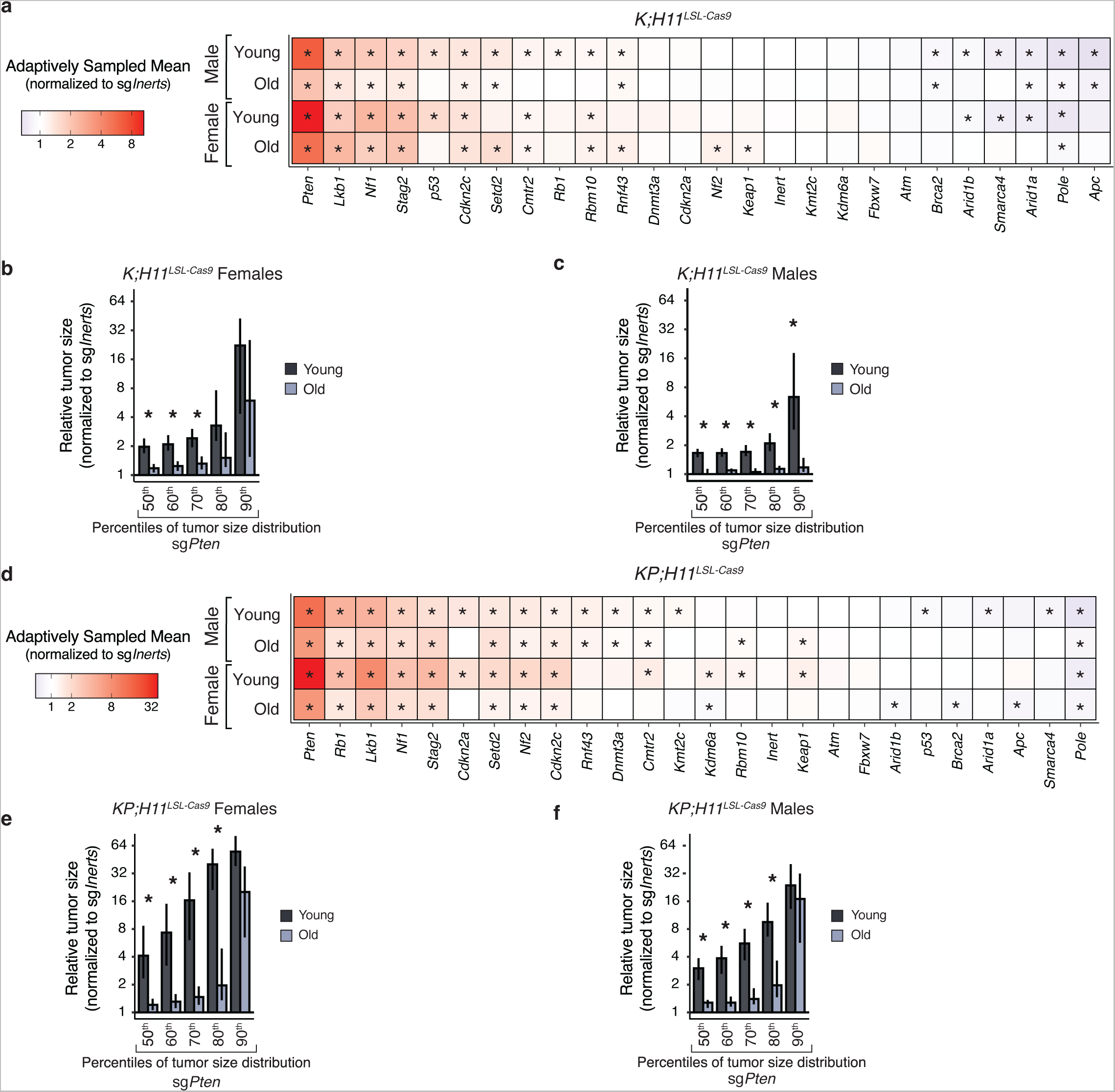
Differential effects of tumor suppressor gene inactivation with age in males and females. **a.** Adaptively sampled mean tumor sizes normalized to sg *Inert* for all tumor genotypes in male(top) and female (bottom) young and old *K;H11^LSL-Cas9^* mice transduced with the Lenti-sg*RNA^Aging^/Cre* pool. Genes are ordered by ASM in young males. Stars denote that gene signicantly impacts ASM relative to sg*Inerts* (two-sided FDR-corrected p<0.05). **b. c.** Adaptively sampled sizes of tumors initiated with sg*Pten* vectors in young and old *K;H11^LSL-Cas9^* female (**b**) and male (**c**) mice transduced with Lenti-sg*RNA^Aging^/Cre* pool at indicated percentiles of the tumor size distribution. Each statistic is normalized to the tumor size at the corresponding percentile of the sg*Inert* distribution. Stars denote a statistically significant difference between young and old (two-sided FDR-adjusted p-value < 0.05). P-values and 95% confidence intervals were calculated using nested bootstrap resampling. **d.** Adaptively sampled mean tumor sizes normalized to sg*Inert* for all tumor genotypes in male(top) and female (bottom) young and old *KP;H11^LSL-Cas9^* mice transduced with the Lenti-sg*RNAAging/Cre* pool. Genes are ordered by ASM in young males. Stars denote that gene signicantly impacts ASM relative to sg*Inerts* (two-sided FDR-corrected p<0.05). **e,f.** Adaptively sampled sizes of tumors initiated with sg*Pten* vectors in young and old *KP;H11^LSL-Cas9^* female (e) and male (f) mice transduced with Lenti-sg*RNA^Ag’ng^/Cre* pool at indicated percentiles of the tumor size distribution. Each statistic is normalized to the tumor size at the corresponding percentile of the sg*Inert* distribution. Stars denote a statistically significant difference between young and old (two-sided FDR-adjusted p-value < 0.05). P-values and 95% confidence intervals were calculated using nested bootstrap resampling.

**Extended Data Fig. 8.**
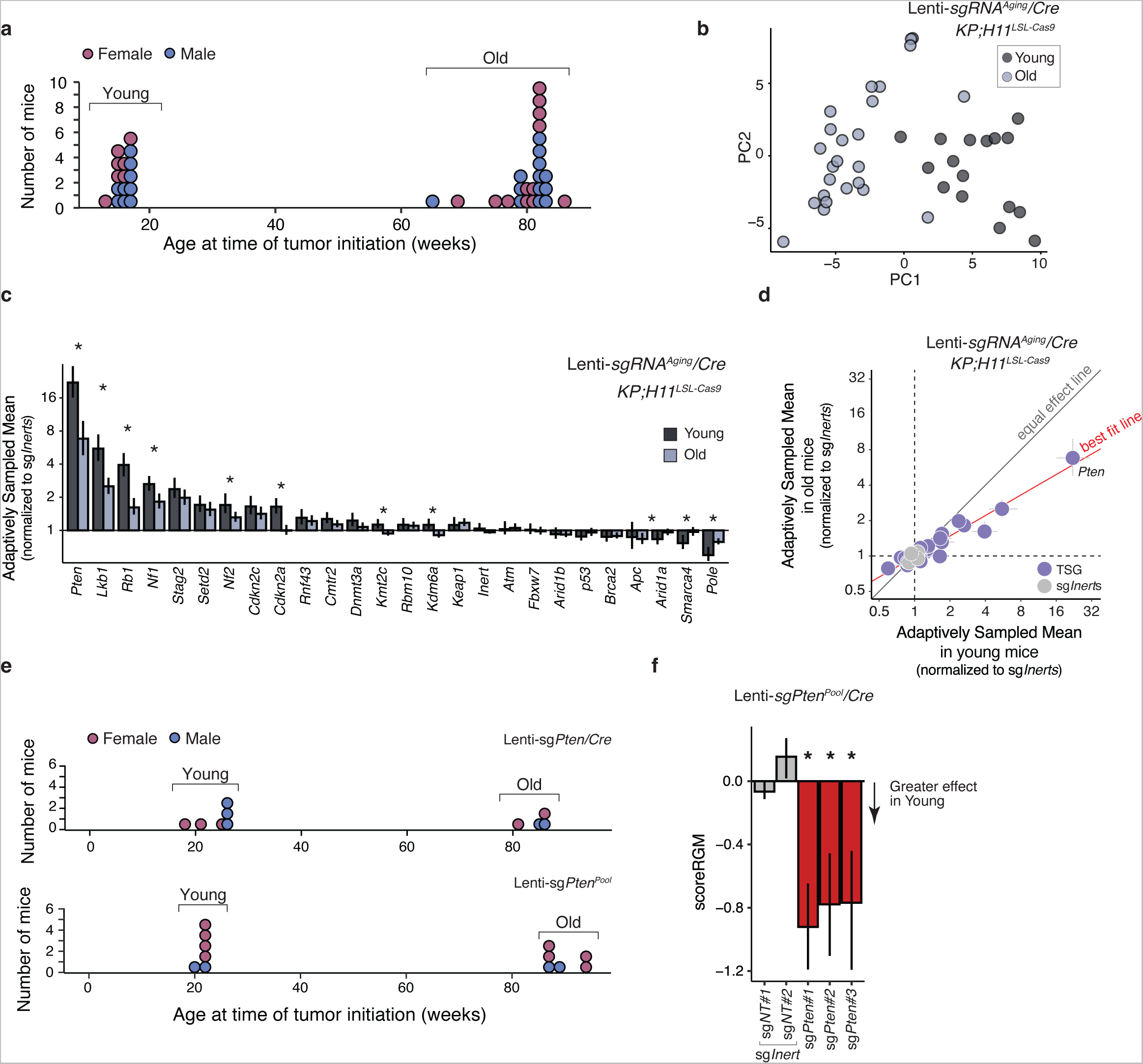
Validation of changes in tumor suppressor function with age. **a.** Ages of *KP;H11^LSL-Cas^9*mice transduced with Lenti-sg*RNAAg’ng/Cre.* Each dot is a mouse; color denotes sex. **b.** Principal components analysis of tumor suppressive effects performed within each mouse reveals separation of young and old *KP;H11^LSL-Cas^9*mice transduced with Lenti-sg*RNAAgng*/Cre. Each dot is a mouse. **c.** Adaptively sampled mean size (ASM) of each tumor genotype normalized to the ASM of sg*/nert*tumors in the young and old *KPHIILSL Cas^9^*mice cohorts transduced with Lenti-sg*RNAAg’ng*/Cre. ASM is a summary metric of tumor fitness that integrates the impact of inactivating each gene on tumor size and number. Line at x=1 indicates no effect on tumorigenesis. P-values and confidence intervals were calculated using nested bootstrap resampling. Genes are ordered by ASM in the young cohort. **d.** Adaptively sampled mean size (ASM) of each tumor genotype normalized to the ASM of sg*/nert* tumors in the young (x-axis) versus old (y-axis) *KP;H11^LSL-Cas9^* mice cohorts transduced with Lenti-sg *RNA^Aging^*/Cre. **e.** Ages of young and old mice transduced with Lenti-sg*Pten/Cre* in Exp #1 (top) and with Lenti-sg*Pten^Pool/^Cre* pool (bottom). Each dot is a mouse; color denotes sex. **f.** Differential effects of each sgRNA in the Lenti-sg*Pten^Pool^Cre* pool with age as quantified by scoreRGM (Li *et al.).* Stars indicate differential effect with age (FDR-adjusted two-sided p-value <0.05). P-values and confidence intervals were calculated using nested bootstrap resampling.

**Extended Data Fig. 9.**
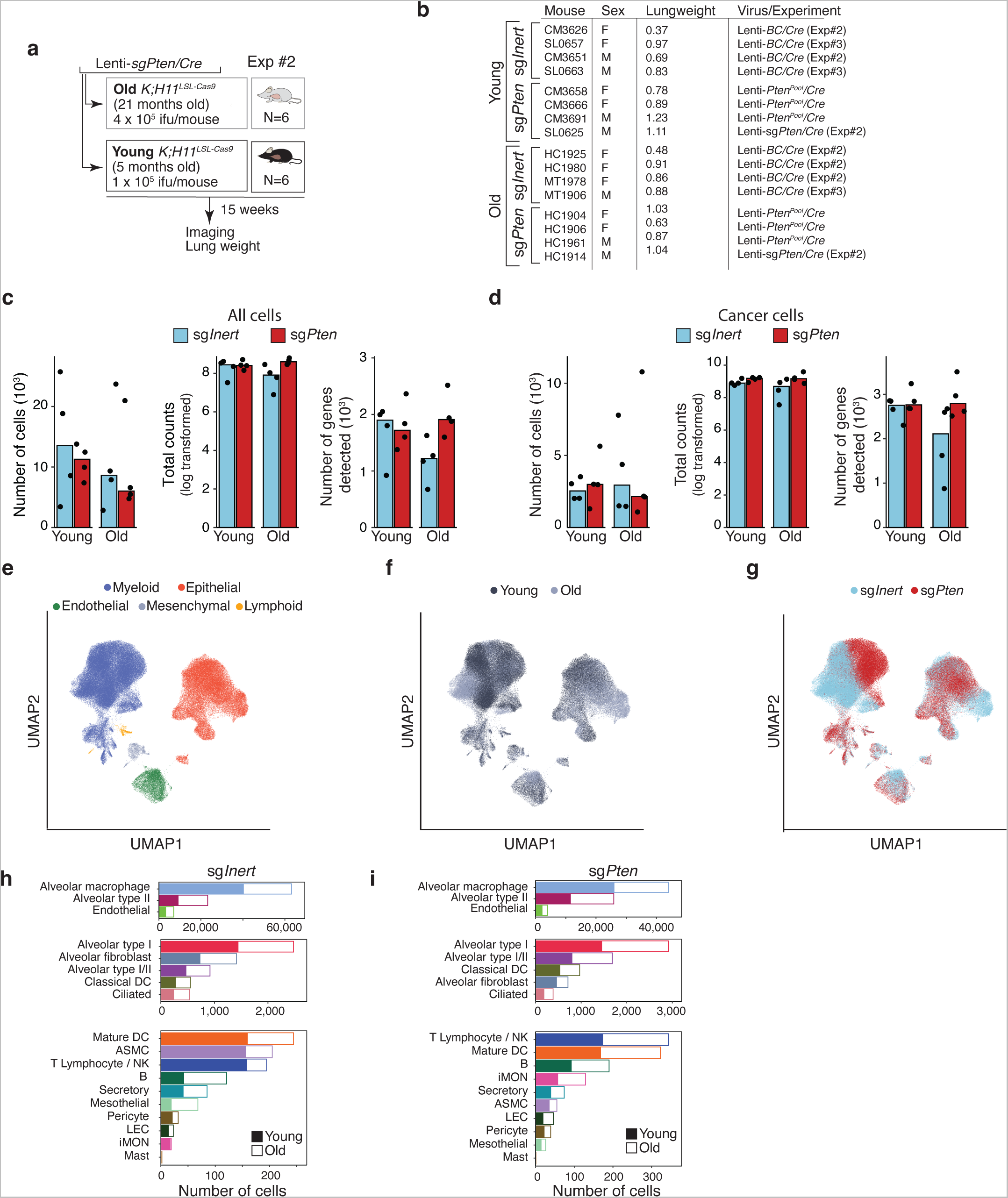
Single-cell RNA sequencing characterizes impacts of aging and *Pten* inactivation on tumor-bearing lungs. **a.** Additional experiment to generate lung samples with *Pten*-deficient tumors. **b.** Table describing characteristics of samples used to generate scRNA-sequencing data. **c.** Total number of cells, median counts per cell, and median number of genes per cell detected per sample after quality control. Each dot is a sample. Tops of bars indicate median values. **d.** Number of cancer cells, median counts per cancer cell, and median number of genes per cancer cell after quality control. Each dot is a sample. Tops of bars indicate median values. **e-g.** Uniform Manifold Approximation and Projection (UMAP) embedding of cells colored by lineage identity (**e**), age (**f**), and tumor genotype (**g**). **h-i.** Counts per cell type by age for sg*Inert* (**h**) and sg*Pten* (**i**) samples. DC: dendritic cells, ASMC: airway smooth muscle cells, iMON: inflammatory monocytes, NK: natural killer cells, LEC: lymphatic endothelial cells.

**Extended Data Fig 10.**
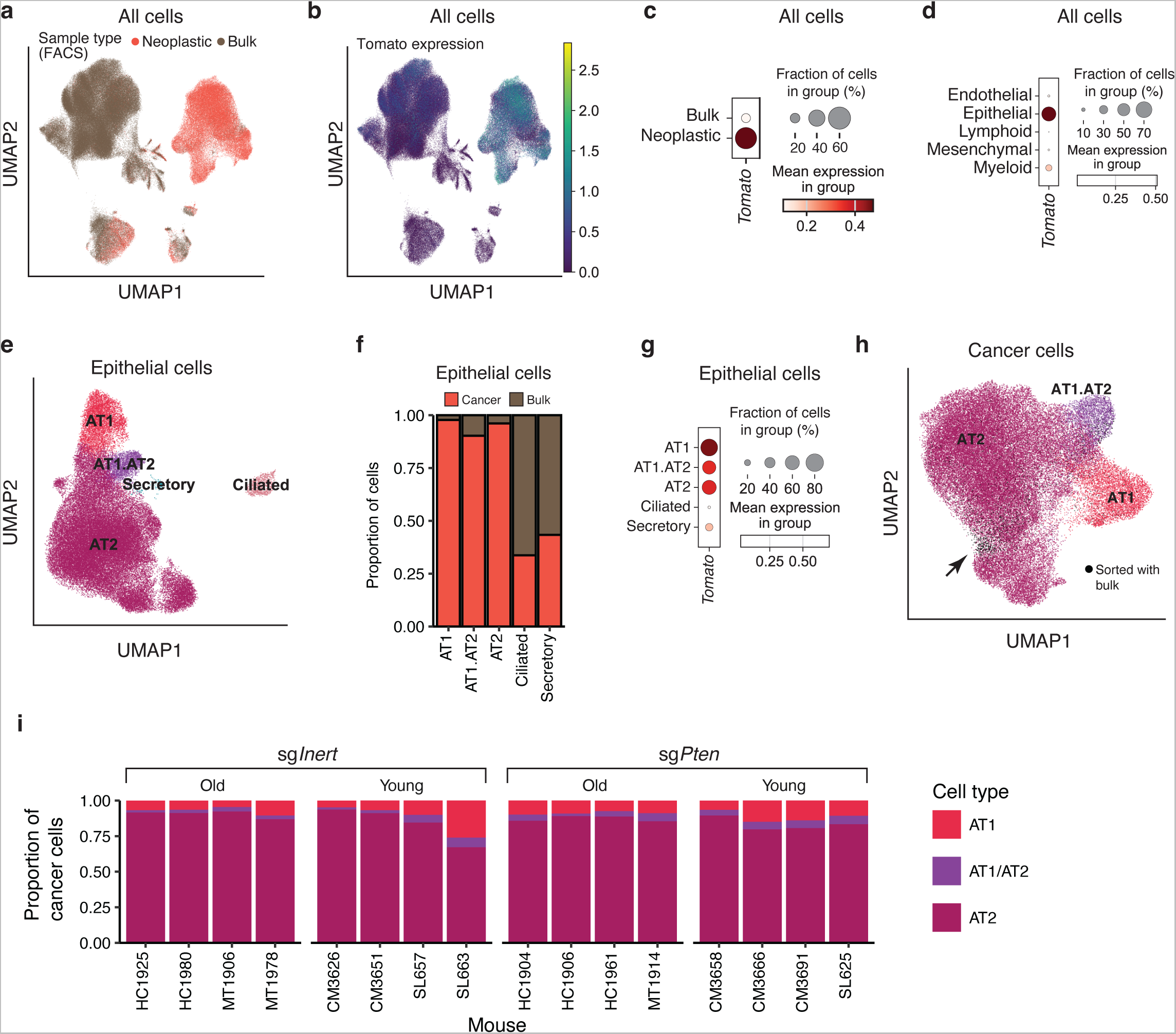
Definition of neoplastic cell populations from single-cell RNA sequencing data. **a.** Uniform Manifold Approximation and Projection (UMAP) embedding of cells colored by whether they were sorted as “neoplastic cells” or “bulk cells” (see **Fig. 4a** and **Methods**). **b-c.** *Tomato* is frequently and highly expressed in cells sorted as neoplastic cells, confirming that sample sorting and preparation worked as expected. **b,** UMAP embedding of cells colored by *Tomato* expression. **c,** Quantification of *Tomato* expression by sample type. **d.** *Tomato* is primarily expressed in cells annotated as epithelial-like cells, consistent with expected phenotype of the lung adenoma and adenocarcinoma cells. **e.** UMAP embedding of epithelial-like cells colored by annotated cell type. **f-g**. The majority of cells annotated as AT1-, AT2-, and AT1/AT2-like cells were sorted as neoplastic cell samples by FACS (**f**) and express *Tomato* (**g**), suggesting that these cell types consist primarily of cancer cells in our dataset. **h.** UMAP embedding of cancer cells (*i.e*. AT1-, AT2- and AT1/AT2-like) cells with the subset of cells sorted into bulk cell samples highlighted in black. Arrow indicates region with highest density of bulk cells. AT1/AT2/AT1.AT2-like cells sorted into bulk cell samples have a higher potential to be untransduced normal lung epithelial cells and were thus excluded from downstream analyses including differential gene expression analysis and age signature scoring (see **Methods**).

**Extended Data Fig 11.**
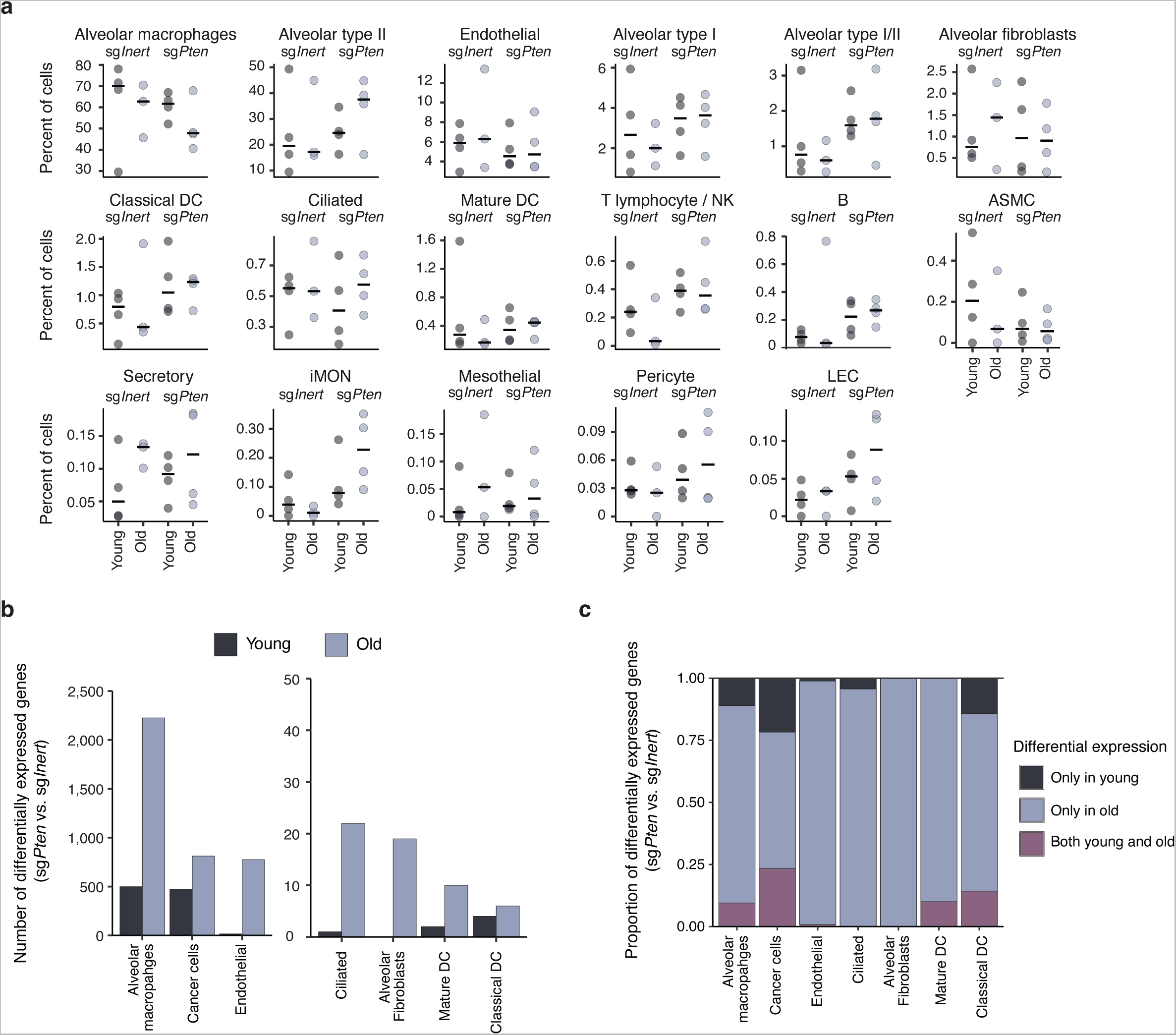
Age shapes the impact of *Pten* inactivation on the tumor microenvironment. **a.** Representation of cell types by age and tumor genotype. Each dot is a mouse; bars indicate median values. Cell types are ordered by overall abundance in the dataset. The impacts of age and tumor genotype on the representation of each cell type were evalulated by two-way ANOVA. No effects were significant after FDR correction. Note that the mouse for which the bulk sample was excluded due to poor viability (MT1906, old sg*Inert*) is excluded. **b.** Number of genes differentially expressed in mice with *Pten*-deficient tumors (Benjamini-Hochberg corrected Wald test p<0.05) in indicated cell types from young and old mice in cell-type resolved pseudobulk differential gene expression analysis. Only cell types with at least two psuedobulked samples (corresponding to at least 10 individual cells and at least 1,000 counts) per age-genotype category were included in the analysis. **c.** Proportion of genes for indicated cell types that that are differentially expressed with the growth of *Pten*-deficient tumors specifically in young mice, specifically in old mice, or in both young and old mice. Genes differentially expressed in both young and old mice were expressed in concordant directions (i.e. overexpressed or underexpressed in both young and old mice with the exception of two genes in alveolar macrophages. DC: dendritic cells.

**Extended Data Fig 12.**
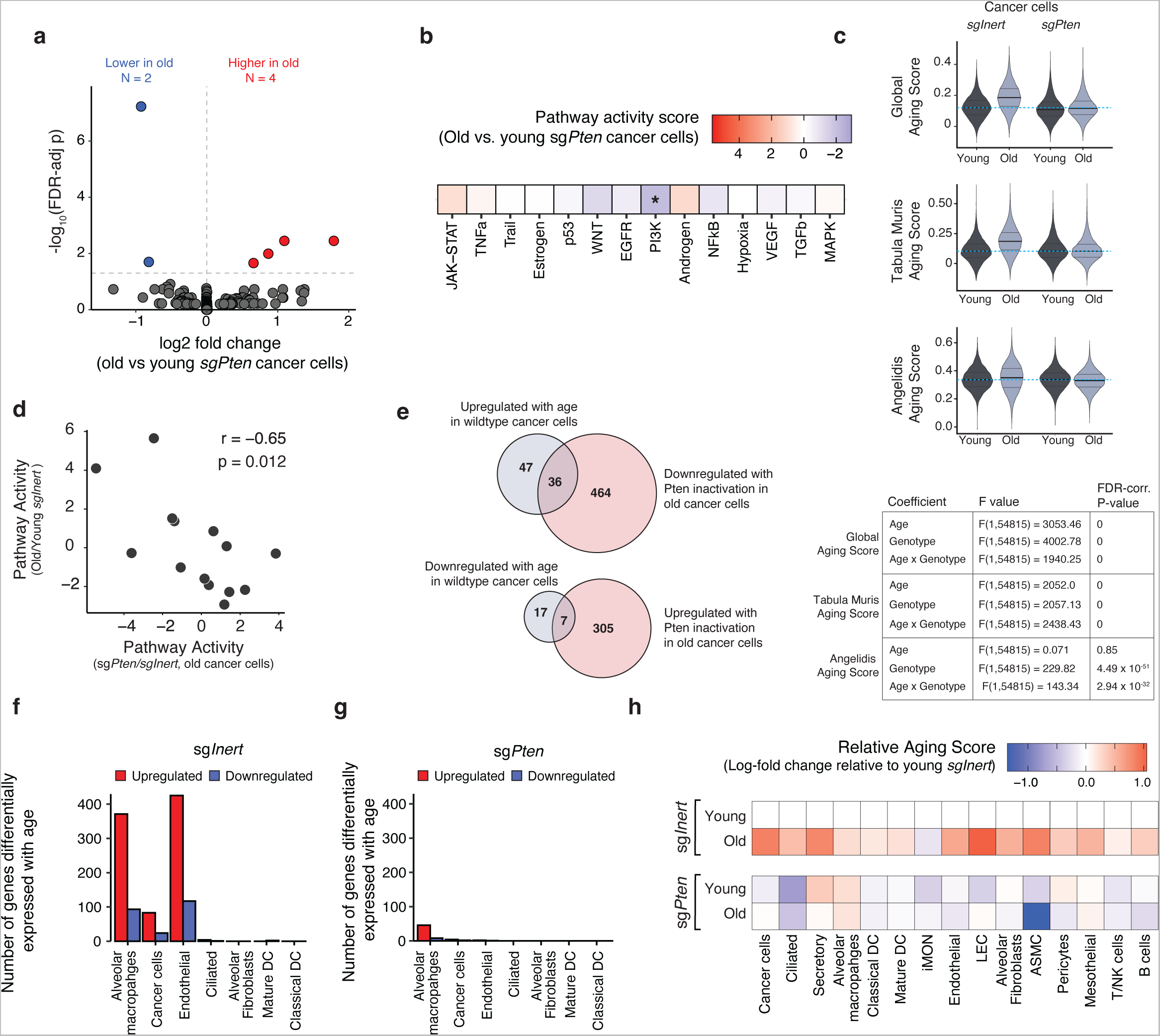
Inactivation of *Pten* decreases molecular phenotypes of aging in cancer cells. **a.** Volcano plot showing changes in gene expression with aging in sg*Pten* cancer cells. Differentially expressed genes (Benjamini-Hochberg corrected Wald test p<0.05) are highlighted in color. **b.** Pathway activity scores comparing old to young cancer cells from sg*Pten* samples. Positive scores (red) indicate that a pathway is more active with age; negative scores (blue) indicate that a pathway is less active with age. Stars denote that a pathway is differentially activated with age (p < 0.05, PROGENy multivariate linear model). Note that the color scale matches the analagous analysis in **Fig. 4f**. **c.** Per-cell scoring of gene expression signatures of normal aging in cancer cells (see **Methods**), stratified by age and genotype (sg*Inert* vs sg*Pten*). Dashed blue line indicates median aging score for young sg*Inert* cells. Results of two-way ANOVA shown in table below. **d.** There is a negative correlation between the effects of *Pten* knockout in old cancer cells and the effects of aging, as measured by pathway activity scores. X-axis corresponds to activity scores in bottom row of **Fig. 4g**; Y-axis coresponds to activity scores in **Fig. 4f**. **e.** Venn diagrams showing overlap beween genes differentially expressed with age and with *Pten* inactivation. Top: roughly 40% of genes that are upregulated with age in wildtype (sg*Inert*) cancer cells and also downregulated with *Pten* inactivation in old cancer cells. Bottom: roughly 30% of the genes that are downregulated with age in wildtype (sg*Inert*) cancer cells and also upregulated with *Pten* inactivation in old cancer cells. Up/down regulation is defined by positive/negative log fold change and Benjamini-Hochberg corrected Wald test p<0.05. **f,g**. Number of genes differentially expressed with age in indicated cell types in mice with sg*Inert* tumors (**g**) and sg*Pten* tumors (**h**). Differential expression determined by Benjamini-Hochberg corrected Wald test p<0.05. Red: upregulated in old. Blue: downregulated in old. **h.** Transcriptomic ages of cells from young and old mice with sg*Inert* and sg*Pten* tumors relative to cells from young mice with sg*Inert* tumors. Cells were individually scored for Global Aging signature (see **Methods**). Shading reflects log-fold change between median age score for each age-genotype grouping and median age score of young-sg*Inert* cells.

